# Blockade of TREM2 ameliorates pulmonary inflammation and fibrosis by modulating sphingolipid metabolism

**DOI:** 10.1101/2024.02.16.580779

**Authors:** Xueqing Gu, Hanyujie Kang, Siyu Cao, Zhaohui Tong, Nan Song

## Abstract

Pulmonary fibrosis is a chronic interstitial lung disease involving systemic inflammation and abnormal collagen deposition. Dysregulations in lipid metabolism, such as macrophage-dependent lipid catabolism, have been recognized as critical factors for the development of pulmonary fibrosis. However, little is known about the signaling pathways involved and the key regulators. Here we found that triggering receptor expressed on myeloid cells 2 (TREM2) plays a pivotal role in regulating the lipid handling capacities of pulmonary macrophages and triggering fibrosis. By integrating analysis of single-cell and bulk RNA sequencing data from patients and mice with pulmonary fibrosis, we revealed that pulmonary macrophages consist of heterogeneous populations with distinct pro-fibrotic properties, and found that both sphingolipid metabolism and the expression of chemotaxis-related genes are elevated in fibrotic lungs. TREM2, a sensor recognizing multiple lipid species, is specifically upregulated in a subset of monocyte-derived macrophages. Blockade of TREM2 by gene knock-out or soluble TREM2 administration can both attenuate bleomycin-induced pulmonary fibrosis. By utilizing scRNA Seq and lipidomics, we found that *Trem2* deficiency downregulates the synthesis of various sphingomyelins, and inhibits the expression of chemokines such as *Ccl2*. Together, our findings not only reveal the alterations in lipidomic profiles and the atlas of pulmonary macrophages during pulmonary fibrosis, but also suggest that targeting TREM2, the crucial regulator affecting both pulmonary sphingolipid metabolism and the chemokines secretion, can benefit pulmonary fibrosis patients in the future.

## 1. Introduction

Pulmonary fibrosis (PF) is a chronic, irreversible lung disease characterized by the deposition of extracellular matrix (ECM), alveolar architecture destruction, which ultimately leads to respiratory failure and death[1, 2]. PF encompasses a wide spectrum of chronic interstitial lung diseases such as idiopathic pulmonary fibrosis (IPF), connective tissue disease-associated ILD (CTD-ILD) as well as organizing pneumonia (OP)[3, 4]. Despite decades of research, the current understanding of different etiologies of pulmonary fibrosis remains unclear. Activated fibroblasts are still the primary target for therapeutic interventions, and very few effective therapies have been discovered[1, 5]. According to previous studies, inhibition of pro-inflammatory signals or additional administration of anti-inflammation agent may also be effective therapeutic strategies [6–8]. Since inflammation may serve as a precursor of fibrosis, especially in the scenario of infection and autoimmune diseases, a series of immune cells have emerged as potential therapeutic targets to ameliorate the progression of fibrosis [5, 8, 9].

Among the immune cells, macrophages have been considered as a key player in triggering pulmonary fibrosis, via the secretion of pro-fibrotic cytokines such as TGF-β and PDGF-AA[10, 11]. However, pulmonary macrophages are functional heterogeneity. Recent advances in single-cell RNA sequencing technology have revealed some fibrosis-associated macrophages populations. Joshi et al. and Misharin et al. reported that monocyte-derived alveolar macrophages can interact with fibroblast, and promote the formation of an essential fibrotic niche[12, 13], while Lyve1^hi^MHCII^lo^ interstitial macrophages alleviate pulmonary fibrosis, as indicated by Chakarov et al.[14]. PRM-151, a modified recombinant human pentraxin 2, which binds with Fcγ receptors and inhibits the differentiation of monocytes into profibrotic macrophages, can improve lung function in IPF patients in a clinical study [15]. These observations suggest that treating specific subset(s) of pulmonary macrophages will benefit the resolution of diseases, and more attention should be paid to identifying the profibrotic macrophage subsets and the critical mechanisms underlying the disease progression.

Dysregulation of lipid metabolism has been characterized as a key feature in pulmonary fibrosis, and lipidomics and metabolomics studies have found that multiple lipid metabolism disorders can be observed in the plasma of both IPF and silicosis patients[16–18]. A series of lipid-derived metabolites, such as ox-LDL, can control the pro-fibrotic response of alveolar macrophages[19]. Ablation of CD36, a scavenger receptor for oxidized phospholipids, alleviates the progression of fibrosis in bleomycin model, probably due to the decreased expression of profibrotic cytokine[20]. Another lipid sensor protein[21], triggering receptor expressed on myeloid cells 2 (TREM2), which is mainly expressed in macrophages, has also been found to be upregulated in both PF patients and lung fibrotic mouse model[13]. However, the role of TREM2 in regulating pulmonary fibrosis is still under debate.

Here, by integrating analysis of single-cell and bulk RNA sequencing in both patients and mice, we confirmed that *TREM2* is upregulated in a specific macrophage subset of the fibrotic lung. Genetic knockout and pharmacological blockade of TREM2 both alleviate bleomycin-induced pulmonary fibrosis, probably due to affecting the sphingolipid metabolism and the chemokine secretion. Taken together, our data confirm the role of TREM2 in regulating the progression of fibrosis, and suggest that targeting lipid-sensors is a promising therapeutic strategy for treating pulmonary fibrosis.

## 2. Materials and methods

### 2.1 Animals

Male C57BL/6J (8-10 weeks) mice weighing 23-28g were purchased from the Vital River Laboratory Animal Technology Co Ltd (Beijing, China), and *Trem2^-/-^* mice (Sock number: 027197) were obtained from the Jackson Laboratory. All mice were kept in a SPF level animal room with a 12 h light/dark cycle. Both male and female *Trem2^-/-^* mice aged 8-10 weeks were used for experiments. All the experiments were approved by the Capital Medical University Animal Care and Use Committee.

### 2.2 Bleomycin induced pulmonary fibrosis mouse model

Pulmonary fibrosis model was induced by intratracheal instillation of 2.5U/kg bleomycin (Nippon Kayaku, Japan) while control group was given the equal volumes of 0.9% saline. On day 21, mice were euthanized under anesthesia after lung function tests, and lung were immediately collected for further analysis.

### 2.3 In vivo inhibition of TREM2 experiments

All C57BL/6J male mice received intratracheal instillation of 2.5U/kg bleomycin on day 0. On the third day, mice were randomly divided into two groups: (1) Bovine Serum Albumin (BSA) and (2) soluble TREM2 protein (sTREM2). On days 3,6 and 9 following intratracheal instillation bleomycin, mice were intranasal inoculation of 0.1mg/kg BSA or sTREM2 protein (Sino Biological, China) in total volume of 10 μL under anesthesia, respectively. On day 21, mice were euthanized under anesthesia after lung function tests, and lung were immediately collected for further analysis.

### 2.4 Survival analysis

Mice were weighed every week after intratracheal instillation of bleomycin to monitor the survival rate. Once the weight loss was over 25% of initial body weight (d0) after intratracheal instillation of bleomycin, the mice were euthanized.

### 2.5 Lung function tests

On day 21 after bleomycin instillation, mice were anesthetized by intraperitoneal injected with 75 mg/kg of 2% pentobarbital. AniRes2005 animal lung function analytic system (Beijing Bestlab High-Tech, China) was used to test lung function. In brief, mice were tracheostomized and inserted endotracheal intubation at a depth of 5∼10mm, then put in the plethysmographic chamber in a supine position. Expiratory velocity was monitored by a pressure sensor, and changes in lung volume were assessed by measuring pressure changes in the plethysmographic chamber. Static lung compliance (CST) was automatically measured by the system[22].

### 2.6 Histology

The right lower lobe was fixed in 4% paraformaldehyde for at least 24 h at room temperature, then dehydrated, embedded in paraffin, and cut into 5-μm slides. Hematoxylin & eosin (HE) staining and Masson trichrome staining were conducted in standard procedures[23]. Collagen volume fraction (collagen area/area of total field) was calculated via Fiji ImageJ software (https://fiji.sc).

### 2.7 Hydroxyproline assay

Levels of hydroxyproline in left lung tissues were measured using the Hydroxyproline Assay kit (Nanjing Jiancheng Bioengineering Institute, Nanjing, China) according to the manufacturer’s instructions. Hydroxyproline content was presented in microgram per left lobe.

### 2.8 RNA isolation and quantitative real-time PCR

Total RNA was extracted from frozen left middle lung lobe using TransZol Up Plus RNA Kit (TransGen Biotech, Beijing, China) according to the manufacturer’s instructions, then cDNA was synthesized according to the PrimeScript TM II 1st Strand cDNA Synthesis Kit (Takara Bio, Dalian, China). Real-time PCR analysis was performed with the LightCycler 480 SYBR Green I Master (Roche) using the ABI Prism 7500 Sequence Detection System instrument. The data was calculated by using 2^- ΔΔCT^ method with *Gapdh* as the internal reference gene. Primers were as follows:

*Col1α1* 5’- GCTCCTCTTAGGGGCCACT-3’ and 3’-CCACGTCTCACCATTGGGG-5’;

*Trem2 5*’ - CTGGAACCGTCACCATCACTC-3’ and 3’-CGAAACTCGATGACTCCTCGG-5’ ;

*Ccl2* 5’ TTAAAAACCTGGATCGGAACCAA-3’ and 3’ GCATTAGCTTCAGATTTACGGGT-5’;

Ccl*7* 5’- GCTGCTTTCAGCATCCAAGTG-3’ and 3’ CCAGGGACACCGACTACTG-5’;

*Gapdh* 5’- AGGTCGGTGTGAACGGATTTG -3’ and 3’-TGTAGACCATGTAGTTGAGGTCA-5’.

### 2.9 Downloading RNA-Seq data

The RNA-Seq data (GSE150910) [24] contained 103 controls and 103 IPF patients was obtained from the National Center for Biotechnology Information (NCBI) Gene Expression Omnibus (GEO). Single-cell RNA-Seq data of bleomycin-indued pulmonary fibrosis mice were downloaded from the GEO database (GSE131800)[25], and single cell RNA seq data of three healthy controls and three IPF patients were accessed from GEO with accession number GSE128033 [26].

### 2.10 scRNA-Seq assay

For scRNA-Seq, fresh right lung of WT and *Trem2^-/-^* mice was minced into small pieces with scissors and digested in RPMI 1640 culture medium (Gibco) with 10% fetal bovine serum (FBS, Gibco) and 1% P/S(Gibco). A mix of Collagenase II (Thermo Fisher), Dispase II (Solarbio), Collagenase IV (Solarbio), DNase I (Solarbio), was used for dissociation and digestion on shaker at 37℃ with 300 rpm rotation for 30 min. Then the dissociated cells were filtered through a 70 μm Cell-Strainer (BD) and centrifuged at 300g for 5 min at 4 °C. Cells were counted and assessed for viability after lysing of red blood cells.

### 2.11 RNA sequencing, and data processing

Sequencing was performed with Illumina (NovaSeq 6000 System) according to the manufacturer’s instructions (Illumina, San Diego, CA), the process has been well described before [27]. Cell Ranger (v.4.0.0) was applied to generate raw gene expression matrices, using mouse GRCm38 as reference genome.

### 2.12 scRNA-Seq data analysis

Seurat package (4.2.1) in R program (4.1.2) was used to screen out high-quality cells (200 to 5,500 genes, mitochondria content less than 20% (mice) or less then 15% (human), and hemoglobin content less than 5%) for following analysis. NormalizeData function was used to normalize and homogenize the data. To batch correction on sample level, the NormalizeData, FindVariableFeatures and ScaleData functions of the Seurat package were used in combination with the Harmony R package (0.1.1). Principal components analysis (PCA) and clustering were conducted with 20 PCs and a resolution of 0.5 for mice or 18 PCs and a resolution of 0.5 for human. FindAllMarkers or FindMarkers function were conducted to analyze differential expressed genes between clusters. Cells were annotated based on the expressions of well recognized markers.

### 2.13 Pathway Enrichment

ClusterProfiler R package (4.2.2) was used for Gene Ontology (GO) enrichment analysis and Gene Set Enrichment Analysis (GSEA). We used both KEGG and GO database for GSEA analysis. Using scMetabolism R package (0.2.1) for metabolic pathway analysis in scRNA-Seq data[28].

### 2.14 Sphingolipid metabolism and Fibrosis Score calculation

We downloaded all genes in KEGG pathway 00600 to calculate Sphingolipid metabolism Score. 20 genes which reported to be associated with pulmonary fibrosis and found to be upregulated in both IPF patients, bleomycin and asbestos-induced fibrosis model were used as fibrosis associated gene set [13]. Sphingolipid metabolism and Fibrosis Score were calculated via AddModuleScore functions in Seurat package (4.2.1).

### 2.15 Metabolic pathway Gene Set Variation Analysis (GSVA)

We downloaded 1,730 genes in 84 metabolism pathways from KEGG PATHWAY Database, and metabolic pathway gene set enrichment analysis was performed using GSVA R package (1.42.0).

### 2.16 Luminex Assays

WT and *Trem2^-/^*^-^ mice were sacrificed under anesthetization on day 7 after intratracheal instillation of bleomycin. Lungs were lavaged with cold sterile PBS and then centrifuged at 300g for 5 min at 4 °C. Supernatant was collected to measure cytokines and chemokines levels via ProcartaPlex Multiplex Immunoassay (ThermoFisher) based on Luminex platform according to manufacturer’s recommendations.

### 2.17 Lipidomics

BALF was collected from different groups as described above (control and bleomycin-induced fibrotic lungs or BSA and sTREM2 groups). 100 *μL* of sample was used for UHPLC-MS/MS analysis conducted by Novogene Co., Ltd. (Beijing, China), the process has been well described before [29].

### 2.18 Metabolomics data analysis

The raw data were processed via the Compound Discoverer 3.01 (CD3.1, Thermo Fisher) to perform peak alignment, peak picking, and quantitation for each metabolite. After metabolites quantification and annotation, principal component analysis (PCA) and Orthogonal Projections to Latent Structures Discriminant Analysis was performed via MetaboAnalyst (www.metaboanalyst.ca). The metabolites with VIP > 1, P-value< 0.05 and fold change ≥ 1.2 or ≤ 0.833 were considered as differential metabolites.

### 2.19 Statistical analysis

Statistical analyses were performed by GraphPad Prism 9.0, and P values ≤ 0.05 were assigned as significant. An unpaired Student’s t-test with two tailed P value was performed for comparisons between two groups. Data were expressed as mean ± SEM. Kaplan Meier survival curves were analyzed with the log-rank (Mantel-Cox) test via GraphPad Software.

## 3. Results

### 3.1 Sphingolipid metabolism is significantly altered during pulmonary fibrosis

A series of recent studies have suggested a strong association between the dysregulation of lipid metabolism and the development of interstitial lung diseases including the idiopathic pulmonary fibrosis (IPF) [16–18]. To decipher the critical lipid metabolites that contribute to the progression of IPF, we first established the bleomycin (BLM)-induced pulmonary fibrosis mouse model. As described, male C57BL/6J mice were intratracheal instillation of 2.5U/kg bleomycin to induce pulmonary inflammation and subsequent fibrosis, which highly resembles IPF. Consistent with previous reports [30], the survival rates in bleomycin-treated mice are significantly lower than that in control group (Supplementary Fig. 1A). The pulmonary function was further measured at day 21 in survival mice. We confirmed that static lung compliance (CST) of bleomycin-induced fibrosis mice is significantly declined compared with control mice (Supplementary Fig. 1B). To compare collagen content between the two groups, the left lung lobes were individually processed for the hydroxyproline assay. As shown in Supplementary Fig. 1C, the hydroxyproline levels are increased after bleomycin treatment. Both Masson’s trichrome staining and quantitative RT-PCR indicated the increased collagen synthesis in the bleomycin-treated mice (Supplementary Fig. 1D-F). These data demonstrated the bleomycin-induced fibrosis mice can be utilized for further studies.

We next investigate the alterations of lipid metabolism in bleomycin-treated mice. The bronchoalveolar lavage fluid (BALF) from both bleomycin-induced fibrosis mice and control mice were collected 21 days post treatment, and analyzed by the quantitative metabolomic approach. A total of 6 molecular species have been quantified covering lipid categories including sphingolipids, glycerolipids and glycerophospholipids, *etc.* (Supplementary Table 1). Comprehensive analysis showed a significant difference between the two groups (Fig. 1A, Supplementary Fig. 1G). Of note, sphingolipids (SP) - related metabolites, including sphingomyelin (SM), ceramide (Cer), and neutral sphingolipids (Hex1Cer), are significantly upregulated in the BALF of bleomycin-treated mice. In addition, acyl carnitine (AcCa) and lysophosphatidylcholine (LPC) are also increased after bleomycin treatment (Fig. 1A and Supplementary Table 2). In contrast, glycerophosphoglycerols (PG), triradylglycerols (TG), lysobisphosphatidic acid (LBPA) and diglyceride (DG) are mainly increased in control mice. Differential metabolite analysis showed that a series of sphingolipids metabolism-related metabolites including SM(d40:2), SM(d18:1/16:0) and HexCer(d40:1), *etc.* are upregulated in the mouse model of fibrosis (Fig. 1B and C), suggesting that the dysregulation of sphingolipid metabolism contributes to the bleomycin-induced pulmonary fibrosis.

**Fig. 1.**
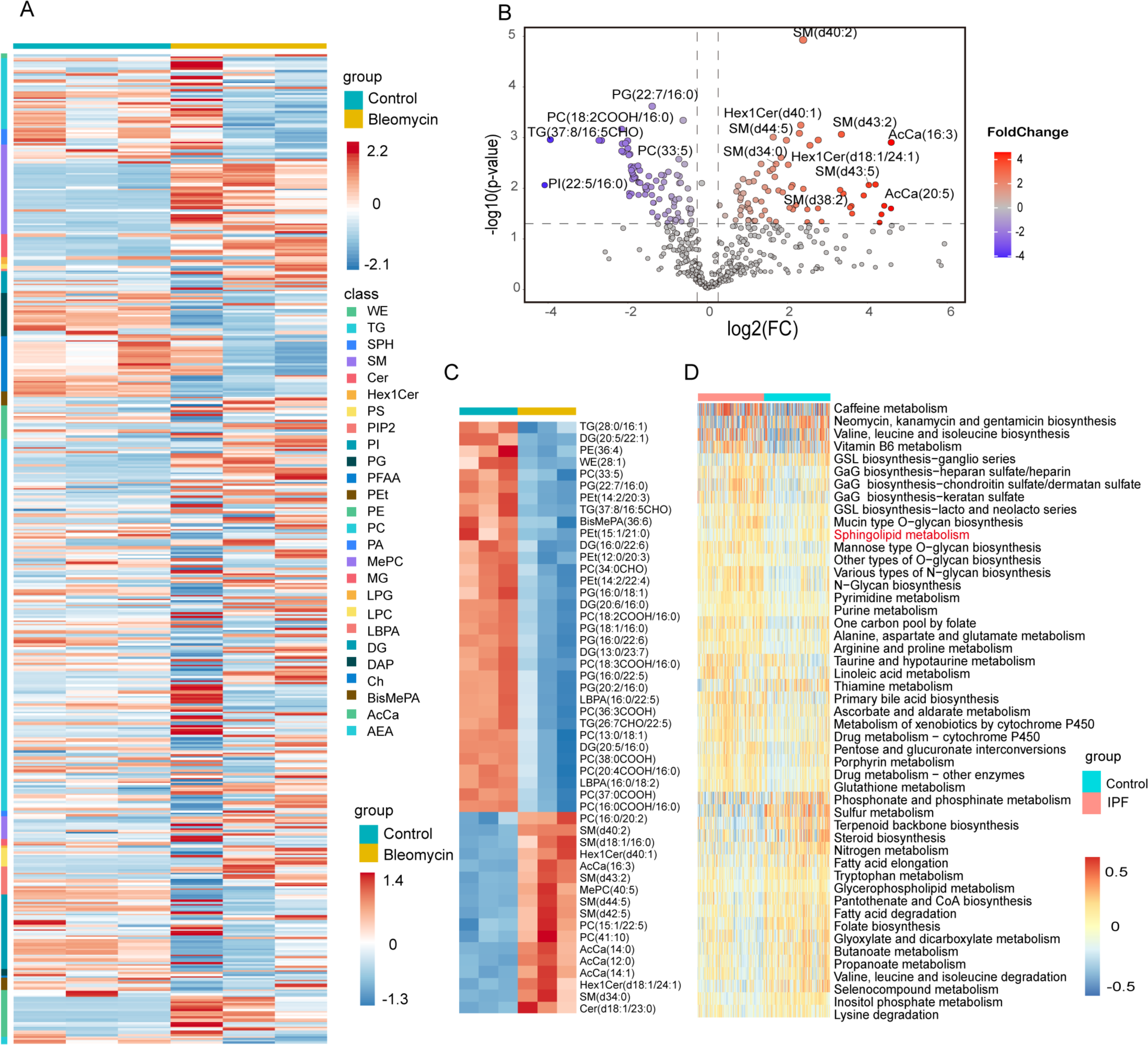
Sphingolipid metabolism is significantly altered during pulmonary fibrosis. (A) Heatmap showing the differentially expressed lipid metabolites subclass between Control and Bleomycin groups (n=3 per group). (B) Volcano plot showing differentially expressed lipid metabolites between Control and Bleomycin groups. Many sphingolipids (SP) -related metabolites were significantly up-regulated in the BALF of bleomycin-treated mice. (C) Heatmap showing the top 50 lipid metabolites in each sample. (D) Heatmap of KEGG metabolic pathway based on GSVA analysis of lung samples from controls and IPF patients. Abbreviations and lipid subclasses are defined in Supplementary Table1.

To further validate the alterations of metabolic pathways in IPF patients, we analyzed the bulk transcriptome sequencing data (GSE150910)[24] from lung samples of IPF patients (n = 103) and control subjects (n = 103) in GEO public platform, based on the major metabolic pathways defined in Kyoto Encyclopedia of Genes and Genomes (KEGG) database (Supplementary Table 3). Principal component analysis (PCA) demonstrated IPF patients and control subjects exhibit distinct metabolic characteristics (Supplementary Fig. 1H). Consistent with previous reports[16, 31, 32], gene set variation analysis (GSVA) demonstrated that the upregulation of *N*-Glycan biosynthesis, glycosaminoglycan biosynthesis, arginine and proline metabolism are associated with pulmonary fibrosis (Fig. 1D). Moreover, the activation of sphingolipid metabolism is also observed in IPF patients. Taken together, these data suggest that aberrant sphingolipid metabolism serves as potential explanations for the progression of pulmonary fibrosis.

### 3.2 TREM2 expression is correlated with dysregulated sphingolipid metabolism during fibrosis

To decipher the potential role of sphingolipid metabolism in pulmonary fibrosis, we then investigated the single-cell RNA Seq data from bleomycin-induced fibrosis mouse model (GSE131800)[25]. After quality control and filtering, 10,400 cells were obtained for the further analysis (Supplementary Fig. 2A). By using differentially expressed genes and canonical markers (Supplementary Fig. 2B), 14 distinct cell types have been defined. As shown in Fig. 2A, the overall number of ATII cells declined obviously, probably due to the treatment of bleomycin, while the proportion of macrophages showed a dramatic increase in the fibrotic lung tissues (Fig. 2A).

**Fig. 2.**
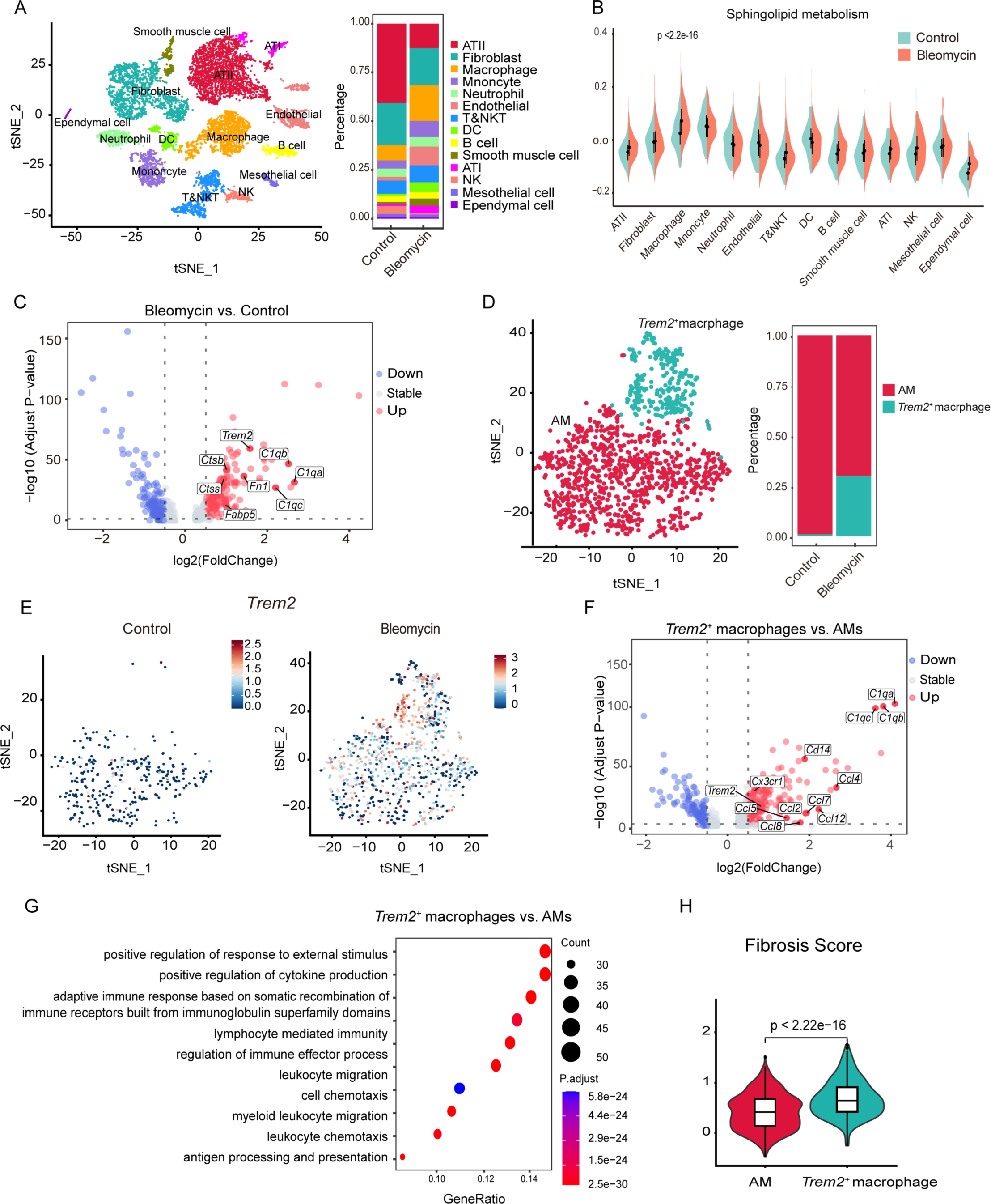
Pro-fibrotic *Trem2*^+^ macrophages are specifically enriched in bleomycin- treated mice. (A) t-SNE plot for all of the 10,400 cells across two experiment conditions (left); the stacked bar plot showing the percentage of each cell type from different groups (right). (B) Violin plot showing Sphingolipid metabolism Score in different cell types between Control and Bleomycin groups. (C) Volcano plot showing differentially expressed genes between macrophages derived from bleomycin-induced fibrotic lungs and control lungs. Negative log2 (FoldChange) indicates downregulation (blue), positive log2 (FoldChange) indicates upregulation (red) in Bleomycin group. (D) t-SNE plot for macrophages (left); the stacked bar plot showing the percentage of each cell type from different groups (right). (E) t-SNE plot showing the expression of *Trem2* in Control (right) and Bleomycin groups (left), respectively. (F) Volcano plot showing differentially expressed genes between *Trem2*^+^ macrophages and AMs. Negative log2 (FoldChange) indicates downregulation (blue), positive log2 (FoldChange) indicates upregulation (red) in *Trem2*^+^ macrophages. (G) Bubble plot showing top 10 GO biological process upregulated in *Trem2*^+^ macrophages. (H) Violin plot showing the gene module scores of fibrosis in AMs and *Trem2*^+^ macrophages.

Since the characteristics of alveolar macrophages (AM) are specialized for lipid metabolism, which is essential for the clearance of pulmonary surfactant. We wondered whether macrophages play a critical role in regulating the sphingolipid metabolism during fibrosis. To confirm this, Sphingolipid metabolism Scores (gene set was from KEGG pathway 00600) were analyzed among the different cell types of lung tissues. As shown in Fig. 2B, the sphingolipid metabolism is predominantly activated in macrophages. Meanwhile, bleomycin treatment further elevated the sphingolipid-related pathways in these cells (Fig. 2B). These suggest that macrophages play an important role in regulating the sphingolipid metabolism during fibrosis, we thus focused on the alterations in gene expression profiles of macrophages during fibrosis. By comparing data of macrophages derived from control and bleomycin-treated groups, we identified a large number of upregulated differentially expressed genes (DEGs) after bleomycin treatment, such as *Fn1, Ctss, Ctsb, etc.*, which are involved in the cell adhesion or ECM-degradation, respectively (Fig. 2C). Most of these genes have been reported to be upregulated in IPF patients [26, 33]. Of note, TREM2, which can bind with sphingolipids[34] and regulate lipid metabolism in various diseases, is also enriched in the macrophages of bleomycin-treated group.

Due to the heterogeneity of pulmonary macrophages, we further subclustered the macrophages (Supplementary Fig. 2C and D), and calculated the fraction of each cell type in both groups (Supplementary Table 4). Surprisingly, we noticed that a population of *Trem2*^+^ macrophages mainly exist in bleomycin-induced fibrosis group (Fig. 2D and E), while AMs are the dominant macrophage population as observed in control mice. Compared with AMs, *Trem2*^+^ macrophages highly express *C1qa, C1qb, C1qc,* and several chemokines including *Ccl2, Ccl4, Ccl7* (Fig. 2F and Supplementary Table 5). Consistently, by GO pathway enrichment analysis, we observed the enrichment of processes such as positive regulation of response to external stimulus, leukocyte migration and cell chemotaxis in *Trem2*^+^ macrophages comparing with AMs (Fig. 2G). Moreover, these *Trem2*^+^ macrophages may be derived from monocytes due to the expression of *Cd14* (Supplementary Fig. 2D). Since monocyte-derived macrophages has been reported as a profibrotic cell subset [18, 19], we then measured the Fibrosis Scores of both AMs and *Trem2*^+^ macrophages, according to the reported Fibrosis gene set [18] (Supplementary Table 6). The Fibrosis Score of *Trem2*^+^ macrophages is significantly higher than that of AMs (Fig. 2H), suggesting a positive correlation between *Trem2* expression and pulmonary fibrosis.

We then validated the upregulation of *TREM2* expression in IPF patients via analyzing RNA Seq dataset (GSE150910) as mentioned above, and found that the *TREM2* expression level is upregulated and associated with sphingolipid metabolism pathway in the IPF patients (Supplementary Fig. 3A and Fig. 3A). To further investigate the role of *TREM2* in IPF patients, a single-cell RNA Seq dataset of lung samples from three healthy controls and three IPF patients (GSE128033) [31] in GEO public platform was analyzed. After quality control and filtering, we obtained 19,012 cells (3,626 cells from control lungs and 15,386 cells from IPF lungs) for further analysis. Unsupervised cluster revealed 19 clusters corresponding to 11 different cell types based on canonical markers (Fig. 3B and Supplementary Fig. 3B and C). *TREM2* expression is significantly higher in IPF patients than healthy controls (Fig. 3C). Since *Trem2* are mainly expressed in monocytes and macrophages, we subclustered all macrophages into seven major clusters, corresponding to three cell types, termed as *FABP4*^+^ macrophages, *GPNMB*^+^ macrophages, and monocytes, respectively (Fig. 3D and Supplementary Fig. 3D-E). Consistent with our previous observations, *TREM2* is mainly expressed in *GPNMB*^+^ macrophages, a specific subpopulation enriched in IPF lungs (Fig. 3E-G).

**Fig. 3.**
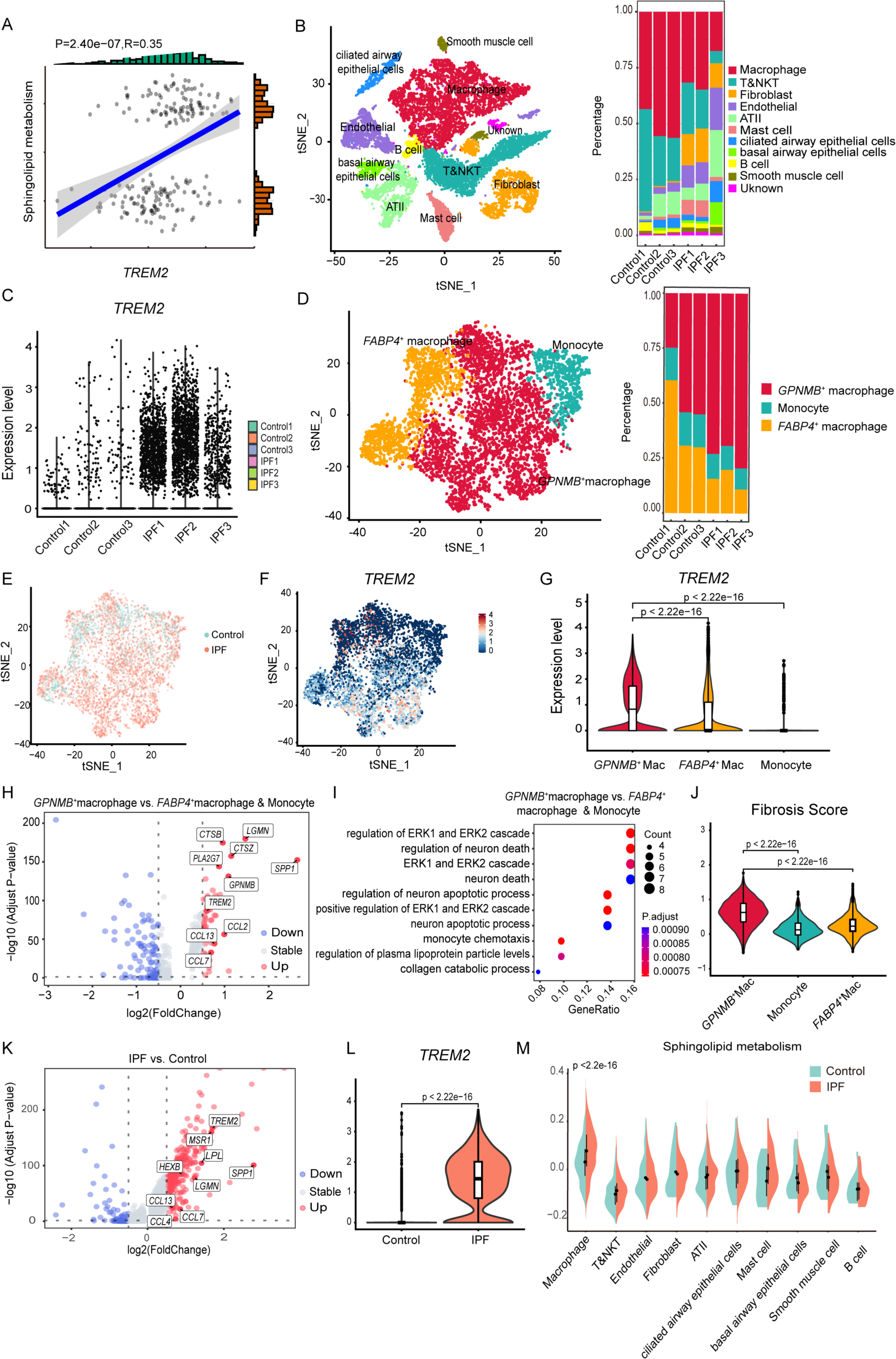
*TREM2* expression is correlated with pulmonary fibrosis and dysregulated sphingolipid metabolism in IPF patients. (A) Scatter plot showing the correlation between *TREM2* expression and sphingolipid metabolism. (B) t-SNE plot for all of the 19,012 cells from healthy controls and IPF patients (left); the stacked bar plot showing the percentage of each cell type from different groups (n=3 per group) (right). (C) Violin plot showing the expression level of *TREM2* in each sample. (D) t-SNE plot for macrophages (left); the stacked bar plot showing the percentage of each cell type from different groups (right). (E) t-SNE plot showing the distribution of macrophages from healthy controls and IPF patients. (F) t-SNE plot showing the expression level of *TREM2*. (G) Violin plot showing the expression level of *TREM2* in each cell type. Mac: macrophage. (H) Volcano plot showing differentially expressed genes between *GPNMB*^+^ macrophages and others. Negative log2 (FoldChange) indicates downregulation (blue), positive log2 (FoldChange) indicates upregulation (red) in *GPNMB*^+^ macrophages. (I) Bubble plot showing top 10 GO biological process upregulated in *GPNMB*^+^ macrophages. (J) Violin plot showing the gene module scores of fibrosis in each cell type. Mac: macrophage. (K) Volcano plot showing differentially expressed genes between *GPNMB*^+^ macrophages from healthy controls and IPF patients. Negative log2 (FoldChange) indicates downregulation (blue), positive log2 (FoldChange) indicates upregulation (red) in IPF patients. (L) Violin plot showing the expression level of *TREM2* in *GPNMB*^+^ macrophages from healthy controls and IPF patients. (M) Violin plot showing Sphingolipid metabolism Score in different cell types between Control and IPF groups.

DEGs analysis was further performed to figure out the differences in gene expression profiles between *GPNMB*^+^ macrophages and other macrophage populations. Besides *TREM2, PLA2G7*, another gene associated with lipid metabolism, is also highly expressed in *GPNMB*^+^ macrophages, consistent with previous report[33]. Similar enrichment can be also observed with regard to ECM-related genes including *CTSB, SPP1* and *LGMN*, as well as chemokines, such as *CCL2, CCL7,* and *CCL13* (Fig. 3H).

The gene expression profiles of *GPNMB*^+^ macrophages resemble that of *Trem2*^+^ macrophages identified in bleomycin-induced fibrosis mice. Functional enrichment analysis of DEGs in *GPNMB*^+^ macrophages showed the activation of ERK1 and ERK2 cascade and monocyte chemotaxis (Fig. 3I). Meanwhile, Fibrosis Score of *GPNMB^+^* macrophages is much higher than those of other two cell types (Fig. 3J). To figure out the potential role of *GPNMB*^+^ macrophages in pulmonary fibrosis, we also compared the gene expression of *GPNMB*^+^ macrophages derived from IPF patients and healthy controls. In addition to the DEGs mentioned before (Fig. 3H), several genes related to lipid metabolism, such as *HEXB, LPL, MSR1* are upregulated in *GPNMB^+^* macrophages of IPF patients (Fig. 3K-L and Supplementary Fig. 3F). Based on the previous findings that aberrant sphingolipid metabolism may be associated with pulmonary fibrosis, we compared Sphingolipid metabolism Score among different cell types. Similar with our previous findings observed in bleomycin-treated mice, Sphingolipid metabolism Score is highest in macrophages, and the sphingolipid metabolism is significantly elevated in macrophages of IPF patients (Fig. 3M). Taken together, our data demonstrated that *Trem2* is predominantly expressed in fibrotic lung tissue, and *Trem2*^+^ macrophages (corresponding to *GPNMB*^+^ macrophages in human patients) may trigger pulmonary fibrosis *via* sensing dysregulated sphingolipid metabolism.

### 3.3 Trem2 deficiency alleviated bleomycin-induced lung fibrosis

To further elucidate the role of *Trem2* in pulmonary fibrosis, we compared the progression of bleomycin-induced pulmonary fibrosis between *Trem2^-/-^* and WT mice. During the 3-week observation period, we found that in *Trem2^-/-^* mice, the mortality rate is much lower (Fig. 4A), and the lung function of CST is improved (Fig. 4B). Meanwhile, hydroxyproline assay showed less collagen accumulation in the lungs of *Trem2*-deficient mice (Fig. 4C), which is further confirmed by Masson’s trichrome staining and quantitative RT-PCR (Fig. 4D-F). Taken together, our data suggest that the expression of *Trem2* is essential for the development of pulmonary fibrosis, and *Trem2* deficiency can alleviated bleomycin-induced fibrosis.

**Fig. 4.**
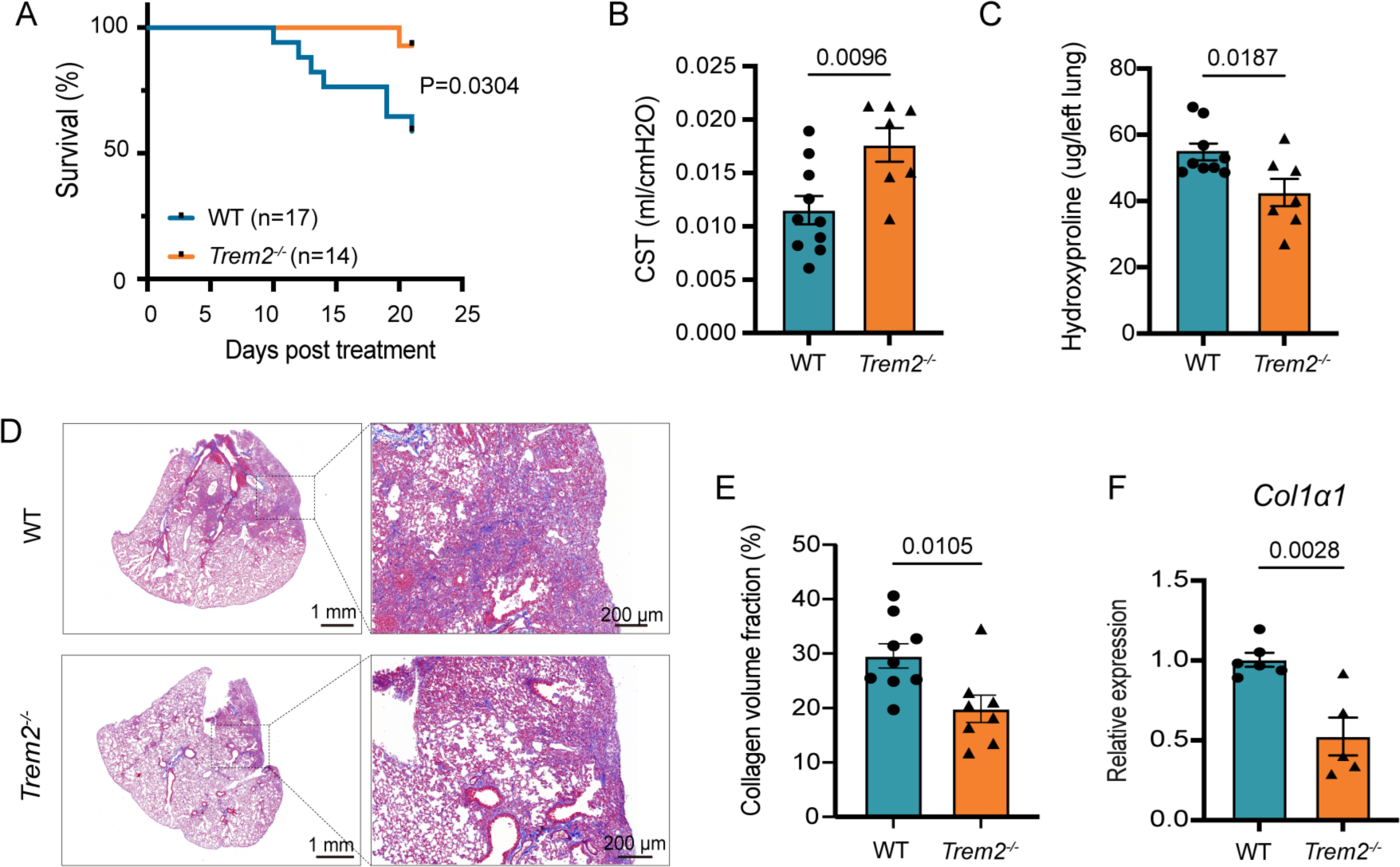
*Trem2* deficiency alleviates bleomycin-induced lung fibrosis. (A) Survival curves of WT and *Trem2^-/-^* mice after bleomycin instillation (n=17 mice in WT group; n=14 mice in *Trem2^-/-^* group). (B) Static lung compliance (CST) of WT and *Trem2^-/-^* mice after bleomycin instillation (n=10 mice in WT group; n=7 mice in *Trem2^-/-^* group). (C) Hydroxyproline contents in lung tissues from WT and *Trem2^-/-^* mice after bleomycin instillation (n=9 mice in WT group; n=7 mice in *Trem2^-/-^* group). (D) Typical images of Masson trichrome staining. Scale bar = 1*mm* (left); Scale bar = 200*μm* (right). (E) Quantitative results of Masson trichrome staining (n=9 mice in WT group; n=8 mice in *Trem2^-/-^* group). (F) The mRNA expression level of *Col1α1* in lung tissues. The data was calculated by using 2^- ΔΔCT^ method with *Gapdh* as the internal reference gene. (n=6 mice in WT group; n=5 mice in *Trem2^-/-^* group). All data are presented as mean±SEM and were analyzed using Student’s t-test.

To clarify the mechanism of *Trem2* deficiency in alleviating bleomycin-induced fibrosis, we collected the lung tissues of both *Trem2^-/-^* mouse and their WT littermate control for single-cell RNA sequencing. After quality control and filtering, 28,813 cells were analyzed, with 15,608 cells from WT and 13,745 cells from *Trem2^-/-^* subjects. Unsupervised clustering revealed 18 cell clusters (Supplementary Fig. 4A), and identified as 12 cell types by using differentially expressed genes and established canonical markers (Fig. 5A and Supplementary Fig. 4B). We identified B cells (*Ms4a1, Cd79a, cd19*), neutrophils (*Csf3r, Retnlg*), T cells and NK cells (*Cd3d, Cd3e, Nkg7, Gzma*), macrophages (*Cd68, Mertk, Adgre1*), monocytes/macrophages (*Ly6c2, Plac8*), DC (*Ccl22, Ccl17*), endothelia cells (*Cdh5, Kdr, Vwf*), fibroblasts (*Pdgfra, Dcn, Col3a1*), myofibroblasts/smooth muscle cells (*Acta2*), ATI (*Hpox, Akap5*) and ATII (*Sftpc*) (Supplementary Fig. 4B) in this dataset. Each cluster is composed of cells derived from both *Trem2^-/-^* and WT mice (Fig. 5B). Of note, *Trem2* is only expressed in some macrophage subpopulations and monocytes/macrophages (Fig. 5C).

**Fig. 5.**
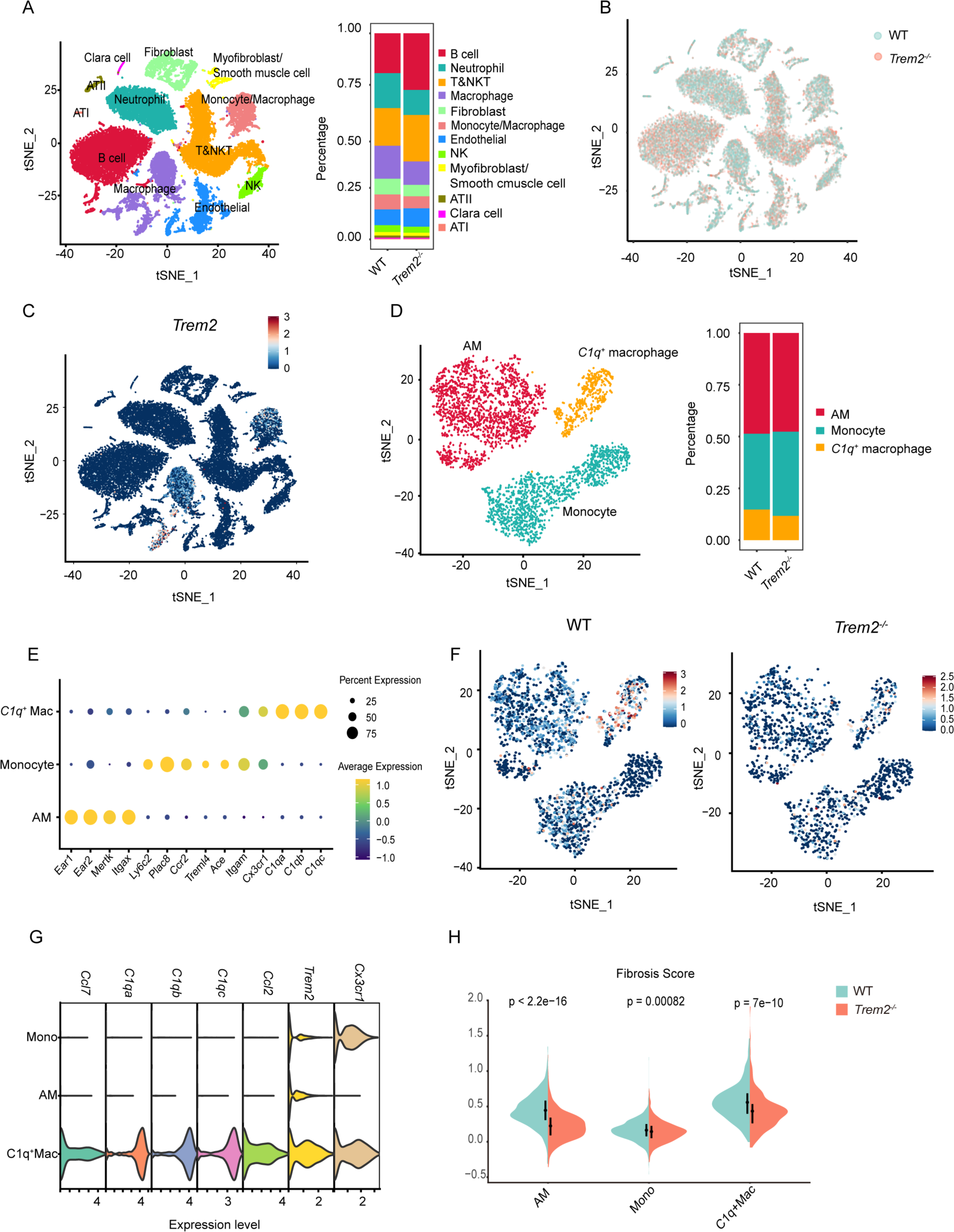
The expression of *Trem2* in the pro-fibrotic *C1q*^+^ macrophages which derived from monocytes in bleomycin-treated mice. (A) t-SNE plot for all of the 28,813 cells from WT and *Trem2^-/-^* groups (left); the stacked bar plot showing the percentage of each cell type from different groups (right). (B) t-SNE plot showing distribution of all cells from WT and *Trem2^-/-^* groups. (C) t-SNE plot showing the expression of *Trem2*. (D) t-SNE plot for macrophages (left); the stacked bar plot showing the percentage of each cell type from different groups (right). (E) Dot plot depicting the average expression levels of canonical cell type-specific marker genes. Mac: macrophage. (F) t-SNE plot showing the expression levels of *Trem2* in WT (left) and *Trem2^-/-^* groups (right), respectively. (G) Violin plot showing selected differently expressed genes in each cell type. (H) Violin plot showing Fibrosis Score between WT and *Trem2^-/-^* groups among three different cell types.

We further subclustered macrophages and distinguished 6 clusters (Supplementary Fig. 4C), and finally identified three cell types indicated as monocytes, alveolar macrophages (AM) and *C1q*^+^ macrophages (Fig. 5D and E). Few significant differences can be observed between the two groups (Fig. 5D and Supplementary Fig. 4D). Similar with *Trem2*^+^ macrophages found in bleomycin-induced fibrosis mice (Fig. 2F), in our datasets, *Trem2* is predominantly expressed in *C1q*^+^ macrophages from WT group (Fig. 5F and G). Meanwhile, *C1q*^+^ macrophages also express relatively high levels of *Cx3cr1* and *Ccr2*, indicating these cells are derived from monocytes (Fig. 5G), similar with previously identified *Trem2*^+^ macrophages [33]. We then compared the Fibrosis Score between WT and *Trem2^-/-^* group among three different cell types as mentioned before (Fig. 2H). The score is highest in *C1q*^+^ macrophages. Moreover, *C1q*^+^ macrophages of WT group has a significantly higher score than those of *Trem2^-/-^* (Fig. 5H).

### 3.4 Trem2 deficiency regulated the sphingolipid metabolism and decreased the levels of sphingomyelin in BALF of bleomycin-treated group

We have showed that dysregulation of sphingolipid metabolism may contribute to the bleomycin-induced pulmonary fibrosis, and sphingolipid signaling pathway is activated in WT compared with *Trem2^-/-^* group (Fig. 6A). Considering the positive correlation between *TREM2* and sphingolipid metabolism (Fig. 3A) in IPF patients, we then speculated that TREM2, as a lipid senser, may modulate sphingolipid metabolism and thus regulate fibrosis. To confirm this hypothesis, we performed KEGG metabolic pathway analysis via the scMetabllism R package [32], and found that the sphingolipid metabolic pathway is upregulated (Fig. 6B) in *C1q*^+^ macrophages compared with others. To clarify the effects of *Trem2* deficiency in sphingolipid metabolism, we investigated sphingolipid metabolism-related genes, and calculated Sphingolipid metabolism Score. Although the score did not show an obvious difference between *Trem2^-/-^* and WT mice (Supplementary Fig. 5A), we found that *Degs1* and *Sgms1*, two key enzymes in sphingolipid metabolism, are both upregulated in WT group (Fig. 6C). These two genes respectively encode dihydroceramide desaturase 1 (DEGS1) and sphingomyelin synthase (SMS), which facilitate the synthesis of ceramide and sphingomyelin (SM). These results suggested that *Trem2* deficiency may affect the synthesis of SM and ceramides, resulting in the dysregulation of sphingolipid signaling pathway.

**Fig. 6.**
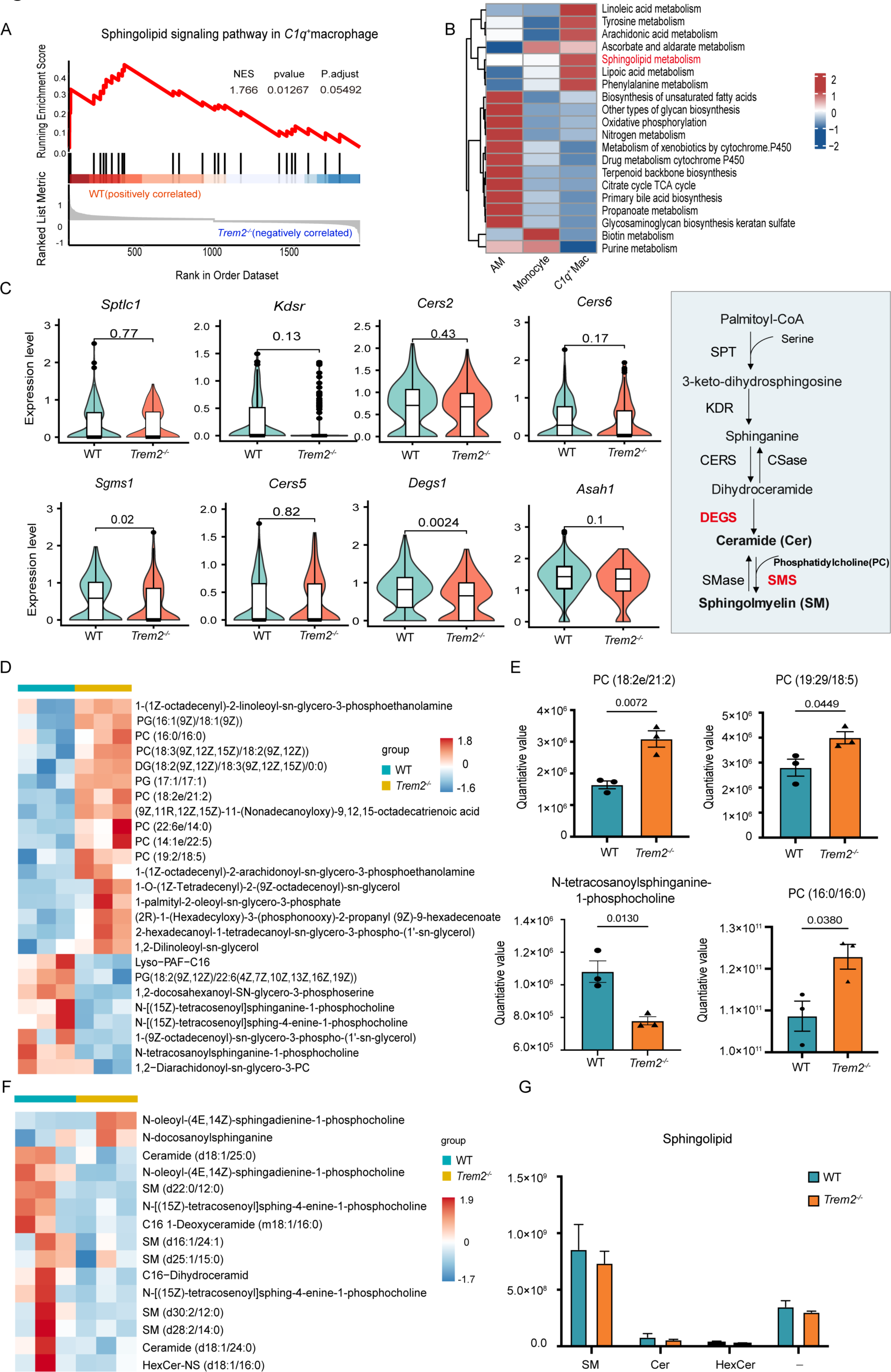
*Trem2* deficiency downregulates the sphingolipid metabolism in bleomycin-treated mice. (A) GSEA analysis identifying the upregulation of Sphingolipid signaling pathway in WT group. (B) Heatmap showing different metabolic pathways in each cell type based on scMetabllism analysis. Mac: macrophage. (C) Violin plots showing the selected genes expression levels between WT and *Trem2^-/-^* groups in *C1q*^+^ macrophages. Schematic diagraph depicting a part of sphingolipid metabolism. SPT: serine palmitoyl transferase; KDR: 3-keto-dihydrosphingosine reductase; CERS: ceramide synthase; CSase: ceramidase; DEGS: dihydroceramide desaturase; SMS: sphingomyelin synthase; SMase: sphingomyelinase. (D) Heatmap showing the top 25 differentially expressed lipid metabolites between WT and *Trem2^-/-^* groups (n=3 per group). (E) Bar plots showing the level of selected differential lipid metabolites in BALF between WT and *Trem2^-/-^* groups (n=3 per group). (F) Heatmap showing the top 15 sphingolipid metabolites between WT and *Trem2^-/-^*groups (n=3 per group). (G) Bar plot showing the sum of different sphingolipid metabolites in each group. All data are presented as mean±SEM and were analyzed using Student’s t-test.

To validate the dysregulation of sphingolipid metabolism causing by *Trem2* deficiency, UHPLC-MS/MS was utilized to detect the alterations of lipid metabolism between *Trem2^- /-^* and WT mice. At day 21 after intratracheal instillation of bleomycin, the BALF of both *Trem2^-/-^* and WT mice was collected for lipidomics analysis. The PCA score plot shows a clear discrimination between metabolic profiles of two groups (Supplementary Fig. 5B) and the heatmap indicated the top 25 differential metabolites (Fig. 6D and Supplementary Table 7). We noticed that *Trem2* deficiency increases the levels of multiple phosphatidylcholines (PC) while decrease the level of N-tetracosanoylsphinganine-1- phosphocholine (SMd18:0/24:0) (Fig. 6E) in the BALF. By investigating the levels of sphingolipid metabolites in our lipidomics, we identified 26 sphingolipid metabolites in total, including 20 sphingomyelin, 4 ceramides, and 2 hexoceramides. Besides, there are 14 sphingolipid metabolites with no corresponding subclassification (-) in the Lipid Subclass (Supplementary Fig. 5C). Although no significant differences has been identified between two groups, multiple sphingolipids including ceramide and sphingomyelin, show a decreasing trend in *Trem2* deficiency group (Fig. 6F and G).

### 3.5 Trem2 deficiency decreased the levels of CCL2 in the BLAF of bleomycin-treated mice

To further investigated the function of *C1q*^+^ macrophages, DEGs analysis was performed between *C1q*^+^ macrophages and other cells (Supplementary Table 8). In addition to some fibrosis-associated genes such as *Lgmn,* many chemokines including *Ccl2, Ccl7, Ccl8, Ccl12* and *Cxcl16* are highly expressed in *C1q*^+^ macrophages (Fig. 7A). GO functional enrichment analysis suggested that the DEGs of *C1q*^+^ macrophages are enriched in pathways related to leukocyte migration, cell chemotaxis and positive regulation of response to external stimulus (Fig. 7B).

**Fig. 7.**
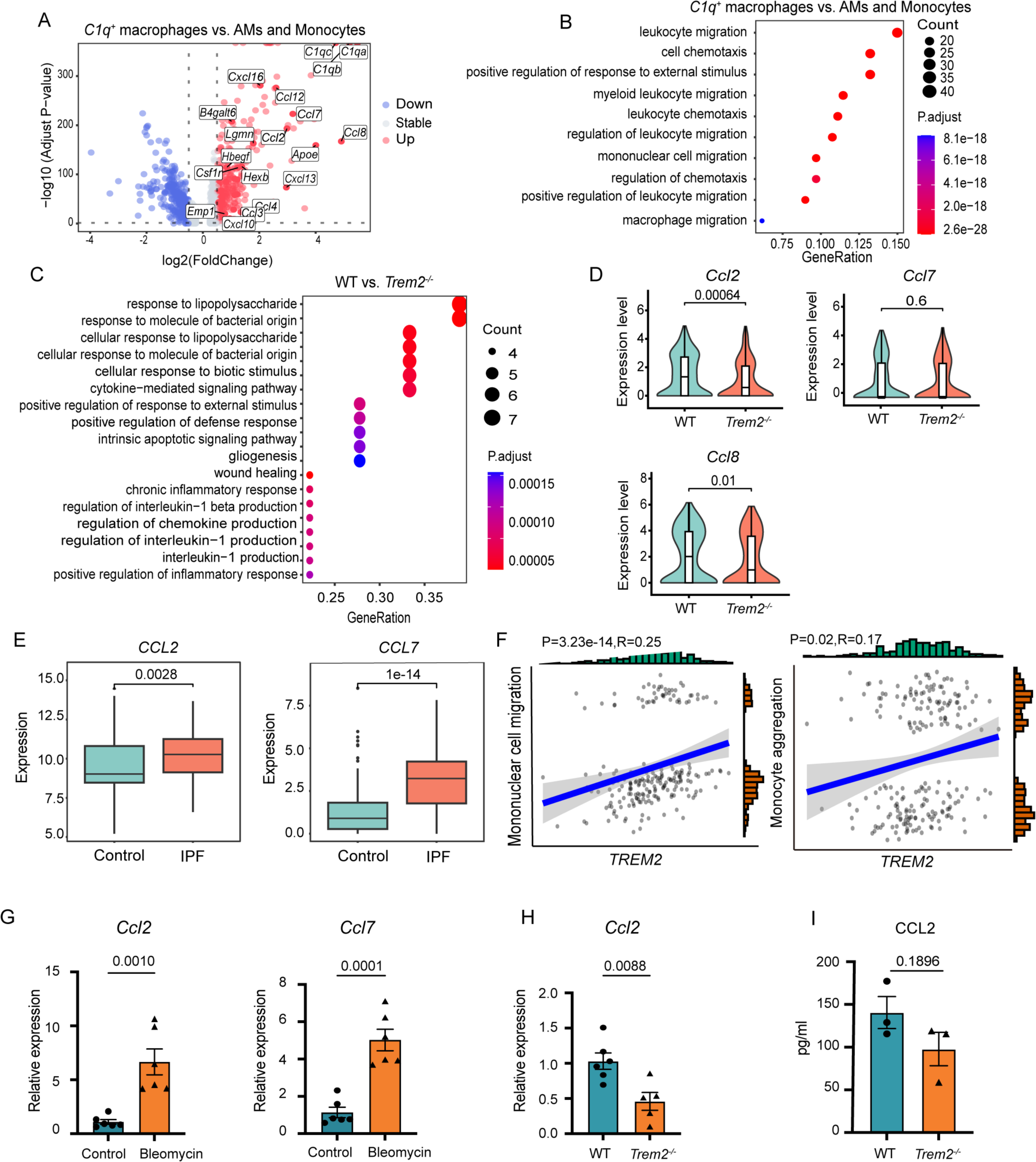
*Trem2* deficiency decreases the levels of chemokines in the bleomycin- treated mice. (A) Volcano plot showing differentially expressed genes between *C1q*^+^ macrophages with others. Negative log2 (FoldChange) indicates downregulation (blue), positive log2 (FoldChange) indicates upregulation (red) in *C1q*^+^ macrophages. (B) Bubble plot showing top 10 GO biological process upregulated in *C1q*^+^ macrophages. (C) Bubble plot showing top 20 GO biological process upregulated in WT compared with *Trem2^-/-^* group in *C1q^+^* macrophages. (D) Violin plots showing the selected genes expression levels in *C1q*^+^ macrophages. (E) Box plots showing *CCL2* and *CCL7* expression levels in RNA Seq data between controls and IPF patients (TPM). (F) Scatter plots showing the correlation between *TREM2* expression and mononuclear cell migration, and the correlation between *TREM2* expression and monocyte aggregation. (G) The mRNA expression levels of *Ccl2, Ccl7* in control and bleomycin-induced mice. The data was calculated by using 2^- ΔΔCT^ method with *Gapdh* as the internal reference gene. (n=6 per group). (H) The mRNA expression level of *Ccl2* in WT and *Trem2^-/-^* groups. The data was calculated by using 2^- ΔΔCT^ method with *Gapdh* as the internal reference gene. (n=6 mice in WT group; n=5 mice in *Trem2^-/-^* group). (I) The level of CCL2 in BALF from WT and *Trem2^-/-^* mice after bleomycin instillation (n=3 per group). All data are presented as mean±SEM and were analyzed using Student’s t-test.

To confirm the effect of *Trem2* deficiency in pulmonary fibrosis, we performed a functional enrichment analysis between *C1q*^+^ macrophages derived from *Trem2^-/-^* and WT groups. Both GO and GSEA functional enrichment analysis suggest that pathways associated to fibrosis, such as response to wounding, is enriched in WT group (Fig. 7C and Supplementary Fig. 6A), while cell chemotaxis, cell migration and inflammatory response are significantly repressed in *C1q*^+^ macrophages of *Trem2^-/-^* group (Fig. 7C and Supplementary Fig. 6B). Moreover, we then examined the chemokines expression level in our data, and confirmed that the expression level of *Ccl2* and *Ccl8* is significantly downregulated in the *Trem2^-/^*^-^ group (Fig. 7D).

Tian et al. reported that CCL2/CCR2 axis can recruit monocyte-derived macrophages and exacerbate profibrotic responses [35]. Considering that *C1q*^+^ macrophages are associated with cell chemotaxis, we thus speculate that *Trem2* deficiency may alleviate fibrosis by downregulating expression of chemokines. We confirmed the upregulation of *CCL2* and *CCL7* in IPF patients via analyzing the RNA Seq data (Fig. 7E). Moreover, *TREM2* shows a positive correlation with mononuclear cell migration, monocyte aggregation and monocyte chemotaxis (Fig. 7F and Supplementary Fig. 6C). In bleomycin-Induced fibrosis mouse model, the expression of *Ccl2* and *Ccl7* is significantly upregulated (Fig. 7G). Moreover, *Trem2* deficiency indeed downregulated *Ccl2* expression (Fig. 7H). ProcartaPlex Multiplex Immunoassay showed that after bleomycin treatment, the secretion of CCL2 showed a decreased trend in the BALF from *Trem2^-/-^* mice, no statistically significant change was evidenced (Fig. 7I and Supplementary Fig. 6D). Taken together, our data showed that *Trem2* deficiency reduce the sphingolipid metabolism as well as the secretion of chemokines in the microenvironment of lungs, which further influence the development of pulmonary fibrosis.

### 3.6 Blockade of TREM2 in vivo ameliorated the bleomycin-induced fibrosis

As *Trem2* deficiency can ameliorate the severity in bleomycin-induced pulmonary fibrosis, we tried to block TREM2 signaling *in vivo* via soluble TREM2 protein (sTREM2), which can competitively bind with TREM2 ligands such as sphingolipids[36]. After intratracheal instillation of bleomycin, mice were randomly divided into two groups, and intranasal inoculation of Bovine Serum Albumin (BSA) or sTREM2 protein at indicated time points, respectively (Fig. 8A). During the 21 days observation period post bleomycin instillation, we found that the mortality rate is slightly decreased after sTREM2 administration (Fig. 8B), and the lung function of CST is also improved by treatment (Fig. 8C). Meanwhile, hydroxyproline assay showed that blockade of TREM2 by sTREM2 leads to a less collagen content, consistent with *Trem2* knockout (Fig. 8D). The results were confirmed by Masson’s trichrome staining and qPCR (Fig. 8E-G). We also examined the *Ccl2* level between BSA and sTREM2-treated groups. The result showed sTREM2 treatment significantly downregulates the expression of *Ccl2* and *Ccl7* (Fig. 8H). Taken together, our data showed that blockade of TREM2 signaling *in vivo* can attenuate the progression of pulmonary fibrosis, improve the survival rate and lung function, and decline the collagen deposition.

**Fig. 8.**
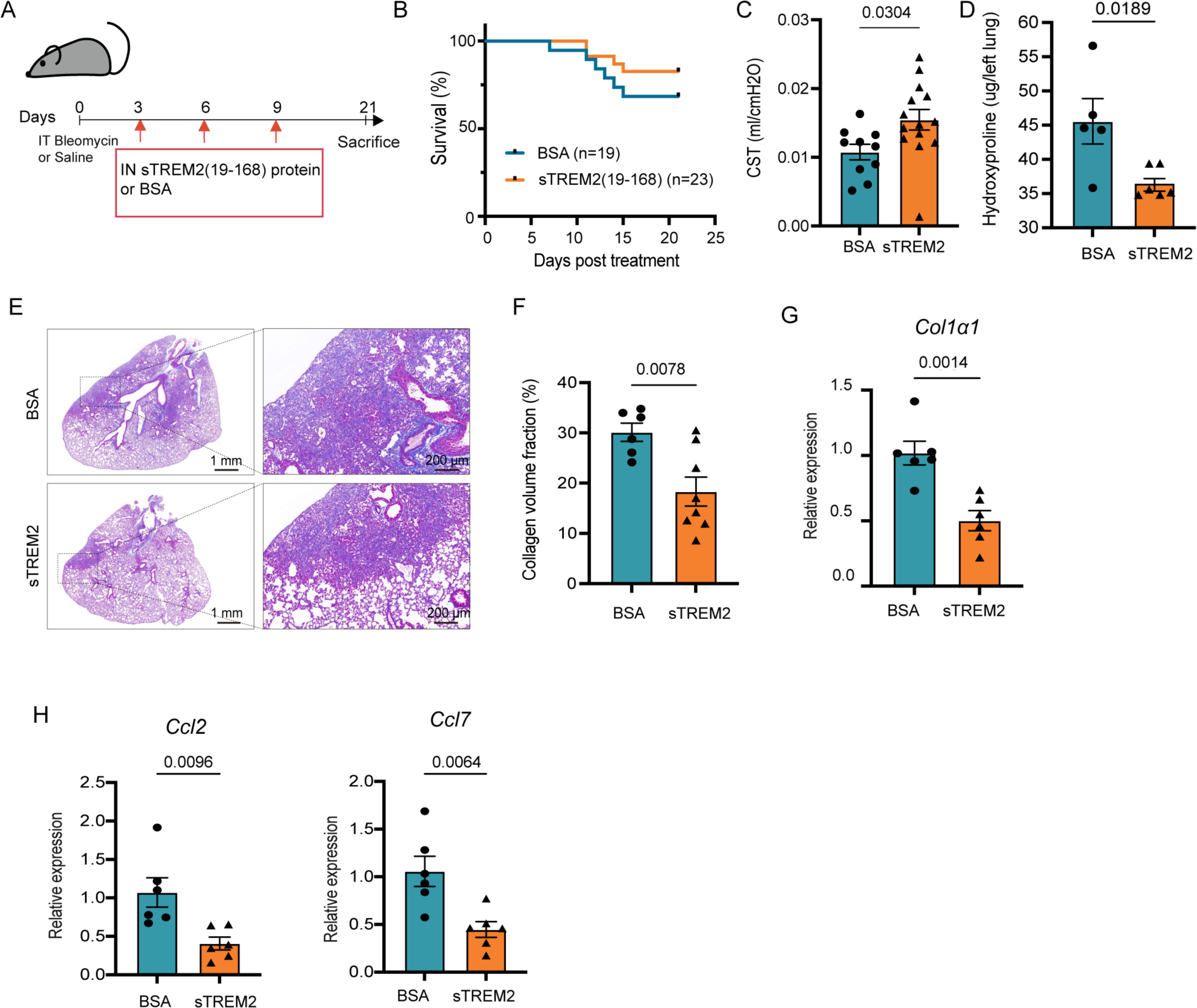
Administration of soluble TREM2 *in vivo* ameliorates the bleomycin-induced fibrosis. (A) Schematic diagram for the treatment schedule and overall study design. BSA: Bovine Serum Albumin. sTREM2: soluble mouse TREM2 protein. (B) Survival curves of bleomycin-induced fibrosis mice after BSA and sTREM2 protein administration respectively (n=19 mice in BSA group; n=23 mice in sTREM2 group). (C) Static lung compliance (CST) of bleomycin-induced fibrosis mice after BSA and sTREM2 protein administration respectively (n=10 mice in BSA group; n=13 mice in sTREM2 group). (D) Hydroxyproline contents in the lung tissues from different groups (n=5 mice in BSA group; n=6 mice in sTREM2 group). (E) Typical images of Masson trichrome staining. Scale bar = 1*mm* (left); Scale bar = 200*μm* (right). (F) Quantitative results of Masson trichrome staining (n=6 mice in BSA group; n=8 mice in sTREM2 group). (G) The mRNA expression level of *Col1α1* in lung tissues. The data was calculated by using 2^- ΔΔCT^ method with *Gapdh* as the internal reference gene (n=6 each group). (H) The mRNA expression levels of *Ccl2* and *Ccl7* in lung tissues. The data was calculated by using 2^- ΔΔCT^ method with *Gapdh* as the internal reference gene (n=6 each group). All data are presented as mean±SEM and were analyzed using Student’s t-test

## 4. Discussion

TREM2 has many biological functions, including induction of phagocytosis, lipid metabolism and anti-inflammation. Phenotypes shift of macrophages triggering by TREM2 may consequence of these processes [37]. In the present study, we reported that TREM2, as a lipid sensor, is upregulated during the progression of pulmonary fibrosis. Blockade of TREM2 signaling by either gene knockout or administration of recombinant soluble TREM2 dramatically attenuates the expression of chemokines such as *Ccl2* in monocyte-derived macrophage, and also decreases the synthesis of sphingolipid metabolites in the pulmonary microenvironments, and further ameliorates experimental pulmonary fibrosis.

Aberrant expression of TRME2 is involved in the development of many diseases such as Alzheimer’s disease and many types of tumors[38–40]. Keren-Shaul reported that TREM2 is essential for the generation of the neurodegenerative microglia, and the lack of TREM2 impairs the transition of microglia from homeostatic to activating state [38, 41, 42]. In Alzheimer’s disease, hematopoietic cell transplantation can reverse TREM2 function via replacing mutant microglia with circulation derived myeloid cells so as to achieve the purpose of treatment [43], suggesting that TREM2 is not only a marker gene for the monocyte-derived cells, but also serves as a critical factor for regulating cell activation. Moreover, Obradovic demonstrated C1Q^+^TREM2^+^APOE^+^ macrophage is a tumor-specific infiltrating macrophage subpopulation, and is associated with disease recurrence in renal carcinoma [40]. Molgora reported that, combination with anti-PD-1, anti-TREM2 monoclonal antibody represses tumor growth [44]. In addition, injection of soluble TREM2 *in vivo* can significantly improve cardiac function in the infarcted heart [45]. Thereby, targeting TREM2 may be an interesting putative therapeutic strategy for treating multiple diseases. Strong enrichment in TREM2 of fibrotic disease has been reported recently [46–48]. A TREM2^+^CD9^+^ subpopulation of macrophages has been found to expand in liver fibrosis, and named as scar-associated macrophages (SAM), which is differentiated from circulating monocytes and profibrotic [46]. We and Luo et al. both found that *TREM2* is upregulated in IPF, and high *TREM2* expression is associated with poor prognosis [49], suggesting a potential role of TREM2 in regulating pulmonary fibrosis. Intriguingly, Hendrikx elucidated that TREM2 can also protect against liver fibrosis, TREM2 deficiency leads to profibrogenic potential and impaired lipid handling in NASH model [50]. Phagocytosis of apoptotic debris induction by TREM2 can prevent endogenous pro-inflammatory signals, leading to restricted inflammation [37]. On the other hand, TREM2-mediated phagocytosis can result in the accumulation of intracellular lipids, which promote the formation of foam cells and the secretion of proinflammatory chemokines [51, 52]. This study indicated that TREM2 may play diverse roles in different settings, probably due to the correlation between the dysregulated lipid metabolism and the progression of diseases.

By intergrading analysis of scRNA Seq and lipidomics of *Trem2^-/-^* and WT mice, we found that *Trem2* deficiency reduces sphingolipid metabolism, especially the synthesis of ceramide and sphingomyelin, and also downregulates the sphingolipid signaling pathway (Fig. 6A and C). Dhami found that sphingolipid metabolism is closely related to the pulmonary fibrosis[53], and sphingolipid signaling pathway may be involved in the development of pulmonary fibrosis [54]. It is hypothesized that ceramide is a signal of lipid overload [55], which may associate with TREM2-dependent phagocytosis. Mayo reported that lactoceramide (LaCer), a metabolite of ceramide synthesized by β-1,4- galactosyltransferase 6 (B4GALT6), can trigger transcriptional programs of *Ccl2* and promote the recruitment of monocytes [56]. Moreover, *B4galt6* was upregulated in *C1q*^+^ macrophages (Fig. 7A). Koh reported that SMS, encode by *Sgms1*, can increase SM levels and decrease PC levels in NASH. Inhibition of *Sgms1* can reduce *Ccl2* expression and alleviate cirrhosis [57]. CCL2 contributes to fibrosis through supporting monocyte/macrophage inflammatory response, angiogenesis and fibroblast recruitment and survival[58–60]. As demonstrated by Suga et al[61]. and Pulito-Cueto et al.[62], the level of CCL2 is upregulated in the serum of IPF and interstitial lung disease (ILD) associated with rheumatoid arthritis (RA) patients. Meanwhile, elevations in CCL2 also have been found in bronchoalveolar lavage[63]. Inhibition of CCL2 signaling can attenuate experimental pulmonary fibrosis [64]. However, in a clinical trial, Carlumab (a humanized monoclonal antibody targeting CCL2) did no provide benefit to IPF patients[60], suggesting that multiple chemokines may be involved in regulating pulmonary fibrosis, and targeting one single chemokine is insufficient for treating these types of diseases. FOXF1, a transcription factor, can regulate the activation of human lung fibroblasts and inhibited macrophages migration via downregulating CCL2, IL-6, TNFα and CXCL1. Bian et al. [65] reported that the transgenic overexpression of FOXF1 can be considered for future therapies in IPF. In our research, *C1q*^+^ macrophages with high expression of *Trem2* is also enrichment in many chemokines, including *Ccl2,* and show strong potential in cell chemotaxis. The level of CCL2 is also downregulated in the BALF of bleomycin-treated *Trem2^-/-^* mice. Moreover, functional enrichment analysis showed blockade of *Trem2* affects the pathways of inflammatory response, leukocyte migration and chronic inflammatory response. Together with our findings that the synthesis of sphingolipid-related metabolites is affected by *Trem2* knockout, these results suggest that sphingolipid metabolism may be involved in regulating fibrosis, probably via triggering the chemotactic and pro-inflammatory phenotype in *C1q*^+^ macrophages.

Specific macrophage subset such as scar-associated macrophages (SAMs) associated with fibrosis in liver, lung and skin has been defined by scRNA Seq technology [46–48]. These cells share a similar expression profiles, including the relatively higher expression of TREM2, C*TSB, CTSL, LGMN, SPP1, PLA2G7, CD63, MMP12, etc.*. Most of these genes are also found in the *C1q*^+^ macrophages identified in our present study (Supplementary Table 8), which also resemble *Trem2*^+^ macrophages (Supplementary Table 5). Moreover, *Trem2* deficiency inhibited the monocyte derived macrophage to fully adopt SAM phenotype in NASH [50]. Since SAM is a profibrotic macrophage subset, *Trem2* may regulate pulmonary fibrosis via affecting the phenotypic transition of monocyte-derived macrophage. However, in our study, we did not find obvious differences in gene expression profiles between *C1q*^+^ macrophages derived from *Trem2^-/-^* and WT groups. However, by re-analysis of AMs of both *Trem2^-/-^* and WT mice, we found a series of differentially expressed genes, and identified a unique cluster characterized by regulation of angiogenesis (data are not shown). Thus, other than triggering chemokine secretion, TREM2 signaling may result in multiple variations during pulmonary fibrosis.

Taken together, our findings demonstrated that *C1q*^+^ macrophages with high expression of *Trem2* play a vital role in promoting fibrosis via regulating sphingolipid metabolism and promoting cell chemotaxis. *Trem2* deficiency can reverse bleomycin-induced pulmonary fibrosis in mice, and in vivo administration of soluble TREM2 recombinant protein to antagonize TREM2 can effectively alleviate fibrosis. These results suggested that TREM2 is promising drug target for IPF therapy.

### CRediT authorship contribution statement

Zhaohui Tong and Nan Song conceived and designed the study. Xueqing Gu and Hanyujie Kang performed the experiments. Xueqing Gu, Hanyujie Kang and Siyu Cao analyzed the data, performed bioinformatic analysis, and drafted the manuscript. Zhaohui Tong and Nan Song revised the final manuscript. All of the authors have approved the final version of the manuscript.

### Data availability statements

The data or code generated during the current study are available from the corresponding author on reasonable request.

## Acknowledgements

This work was supported by the National Natural Science Foundation of China (82070005, 82270009, 82172278); the National Key Research and Development Program of China (2023YFC0872500); the Beijing Natural Science Foundation (JQ22019); the Capital’s Funds for Health Improvement and Research (CFH2022-1-1061); the Beijing Scholars Program (No. 062); the Reform and Development Program of Beijing Institute of Respiratory Medicine.

---

**Supplementary Table 1**. Lipid subclasses definition and abbreviations of lipid metabolites.

**Supplementary Table 2**. Differentially expressed metabolites detected in BALF between bleomycin-induced pulmonary fibrosis and control groups.

**Supplementary Table 3.** KEGG metabolic pathway used for GSVA analysis.

**Supplementary Table 4.** Percentage of macrophages in different groups.

**Supplementary Table 5.** Differentially expressed genes between *Trem2*^+^ macrophages and AMs.

**Supplementary Table 6.** Fibrosis associated genes used for measuring Fibrosis Score.

**Supplementary Table 7.** Differentially expressed metabolites detected in BALF between

*Trem2^-/-^* and WT groups after intratracheal instillation of bleomycin.

**Supplementary Table 8.** Differentially expressed genes between *C1q*^+^ macrophages and other macrophages/monocytes.

Supplementary Fig 1. Aberrant sphingolipid metabolism occurres during pulmonary fibrosis in both bleomycin-induced fibrosis models and IPF patients.

(A) Survival curves of control and bleomycin-induced fibrosis mice (n=12 mice in Control group; n=15 mice in Bleomycin group).

(B) Static lung compliance (CST) of control and bleomycin-induced fibrosis groups (n=10 mice in Control group; n=11 mice in Bleomycin group)

(C) Hydroxyproline contents in the lung tissues from control mice and bleomycin-induced lung fibrosis mice. (n=5 mice in Control group; n=6 mice in Bleomycin group).

(D) Typical images of Masson trichrome staining. Scale bar = 1*mm* (left); Scale bar = 200*μm* (right).

(E) Quantitative results of Masson trichrome staining (n=6 mice in Control group; n=15 mice in Bleomycin group).

(F) The mRNA expression level of *Col1α1* in lung tissues. The data was calculated by using 2^- ΔΔCT^ method with *Gapdh* as the internal reference gene (n=6 per group).

(G) PCA analysis of metabolic profiles of BALF from control and bleomycin mice.

(H) PCA analysis of metabolic pathway profiles of lung samples from healthy controls and IPF patients.

All data are presented as mean±SEM and were analyzed using Student’s t-test

Supplementary Fig 2. Overview of the landscape in bleomycin-induced fibrosis mouse model.

(A) t-SNE plot for all of the 10,400 cells across two experiment conditions colored by clusters.

(B) Dot plot depicting the average expression levels of canonical cell type-specific marker genes.

(C) t-SNE plot for macrophages colored by clusters.

(D) Dot plot depicting the average expression levels of canonical cell type-specific marker genes for macrophages.

Supplementary Fig 3. Pro-fibrotic *GPNMB*^+^ macrophages are specifically enriched in IPF patients.

(A) Box plot showing *TRME2* expression levels between controls and IPF patients (TPM).

(B) t-SNE plot for all of the 19,012 cells across two experiment conditions colored by clusters.

(C) Dot plot depicting the average expression levels of canonical cell type-specific marker genes.

(D) t-SNE plot for macrophages colored by clusters.

(E) Dot plot depicting the average expression levels of canonical cell type-specific marker genes in different macrophages.

(F) Violin plots showing the selected genes expression levels in *GPNMB*^+^ macrophages.

Supplementary Fig 4. Overview of the landscape in WT and Trem2-/- groups after bleomycin instillation.

(A) t-SNE plot for all of the 28,813 cells across two experiment conditions colored by clusters.

(B) Dot plot depicting the average expression levels of canonical cell type-specific marker genes.

(C) t-SNE plot for macrophages colored by clusters.

(D) t-SNE plot for macrophages colored by groups.

Supplementary Fig 5. *Trem2* deficiency alteres sphingolipid metabolism in bleomycin-induced fibrosis.

(A) Violin plot showing the Sphingolipid metabolism Score between two group in *C1q*^+^ macrophages.

(B) PCA analysis of metabolic profiles of BALF from WT and *Trem2^-/-^* mice.

(C) Pie chart showing the composition of sphingolipids (SP) -related metabolites detected in BALF from WT and *Trem2^-/-^* mice after bleomycin instillation.

Supplementary Fig 6. *Trem2* deficiency alters the ability in cell chemotaxis and the expression of CCL2 in bleomycin-induced fibrosis.

(A) GSEA analysis identifying the activation of response to wounding pathway in WT derived *C1q*^+^ macrophages.

(B) GSEA analysis identifying cell chemotaxis, cell migration, chemokine production and regulation of chemokine production pathway were upregulated in WT in *C1q*^+^ macrophages.

(C) Scatter plot showing the correlation between *TREM2* expression and monocyte chemotaxis.

(D) Heatmap showing the measured cytokines and chemokines levels in BALF between WT and *Trem2^-/-^* group after bleomycin

