## Supplemental Figure for "Blockade of TREM2 ameliorates pulmonary inflammation and fibrosis by modulating sphingolipid metabolism"

**Supplementary Fig 1.**

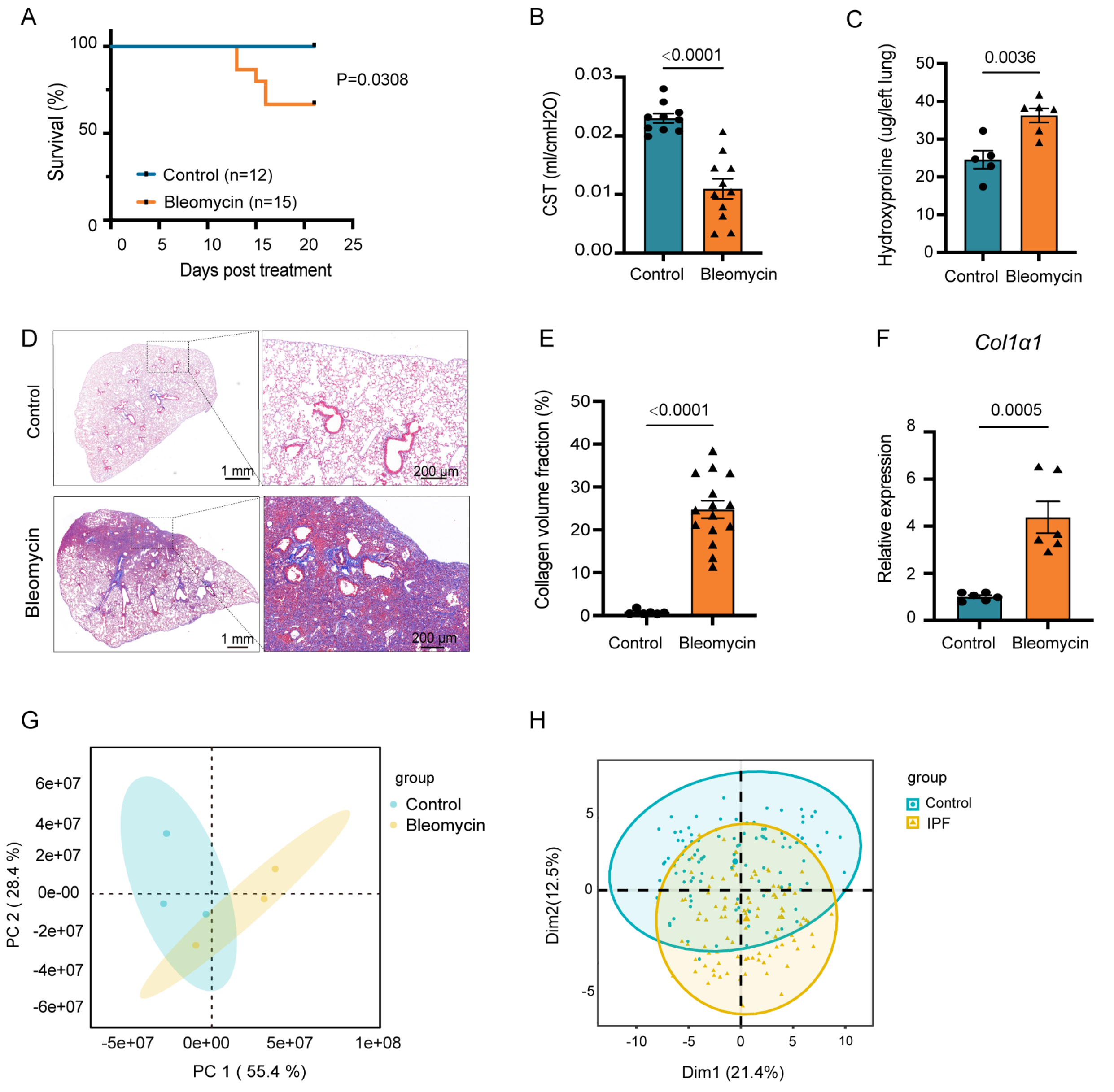

**Supplementary Fig 2.**

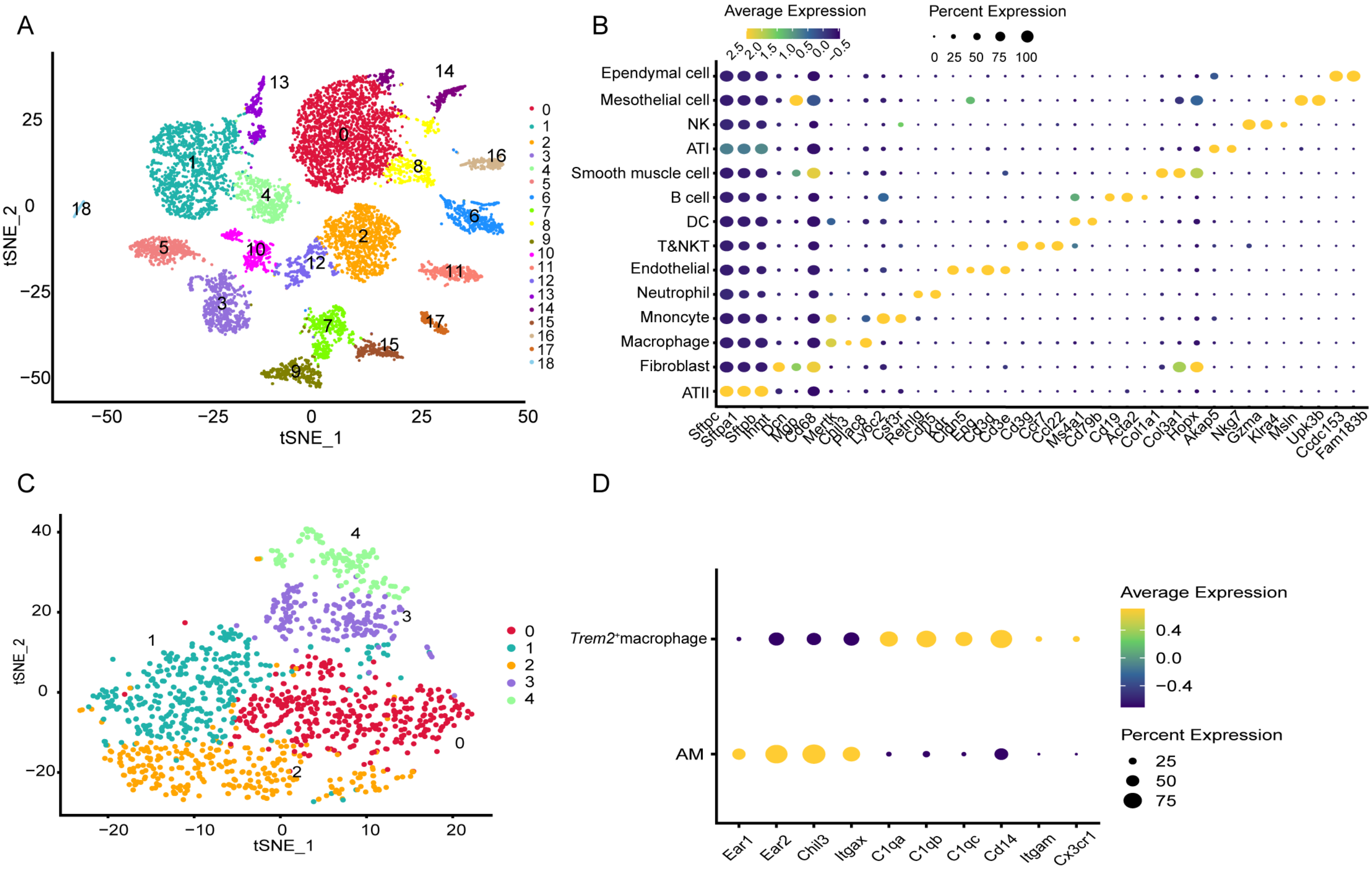

**Supplementary Fig 3.**

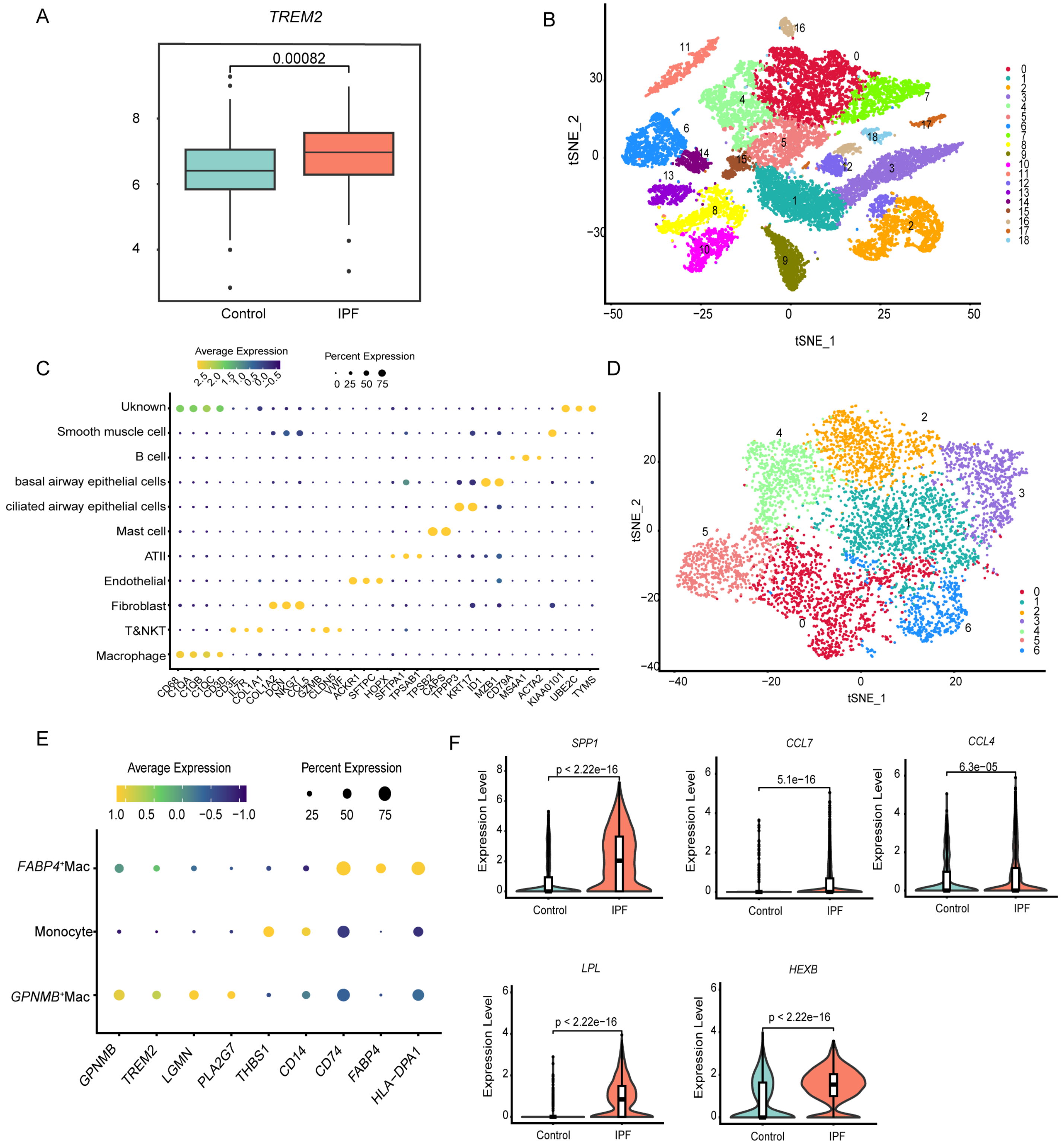

**Supplementary Fig 4.**

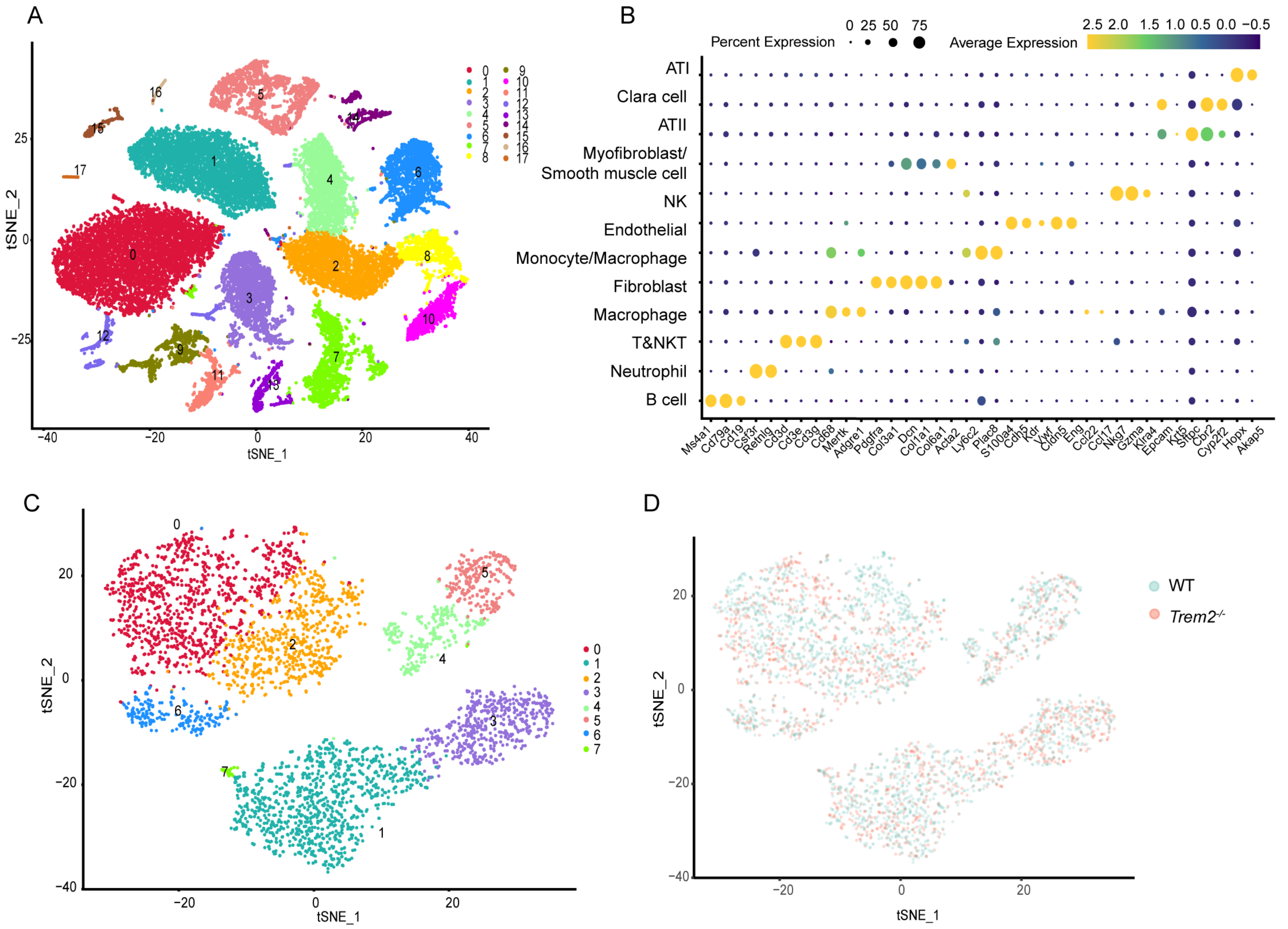

### Supplementary Fig 5.

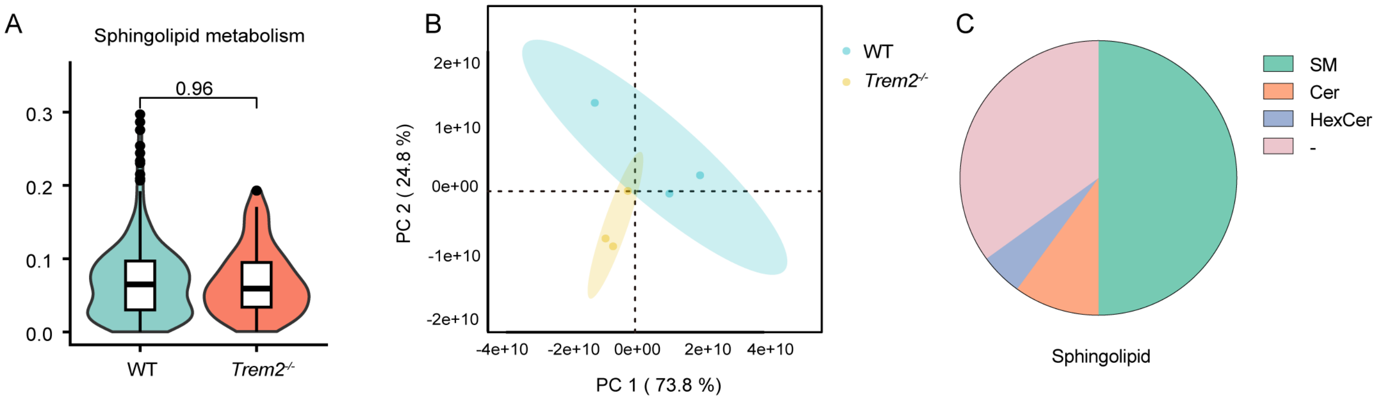

### Supplementary Fig 6.

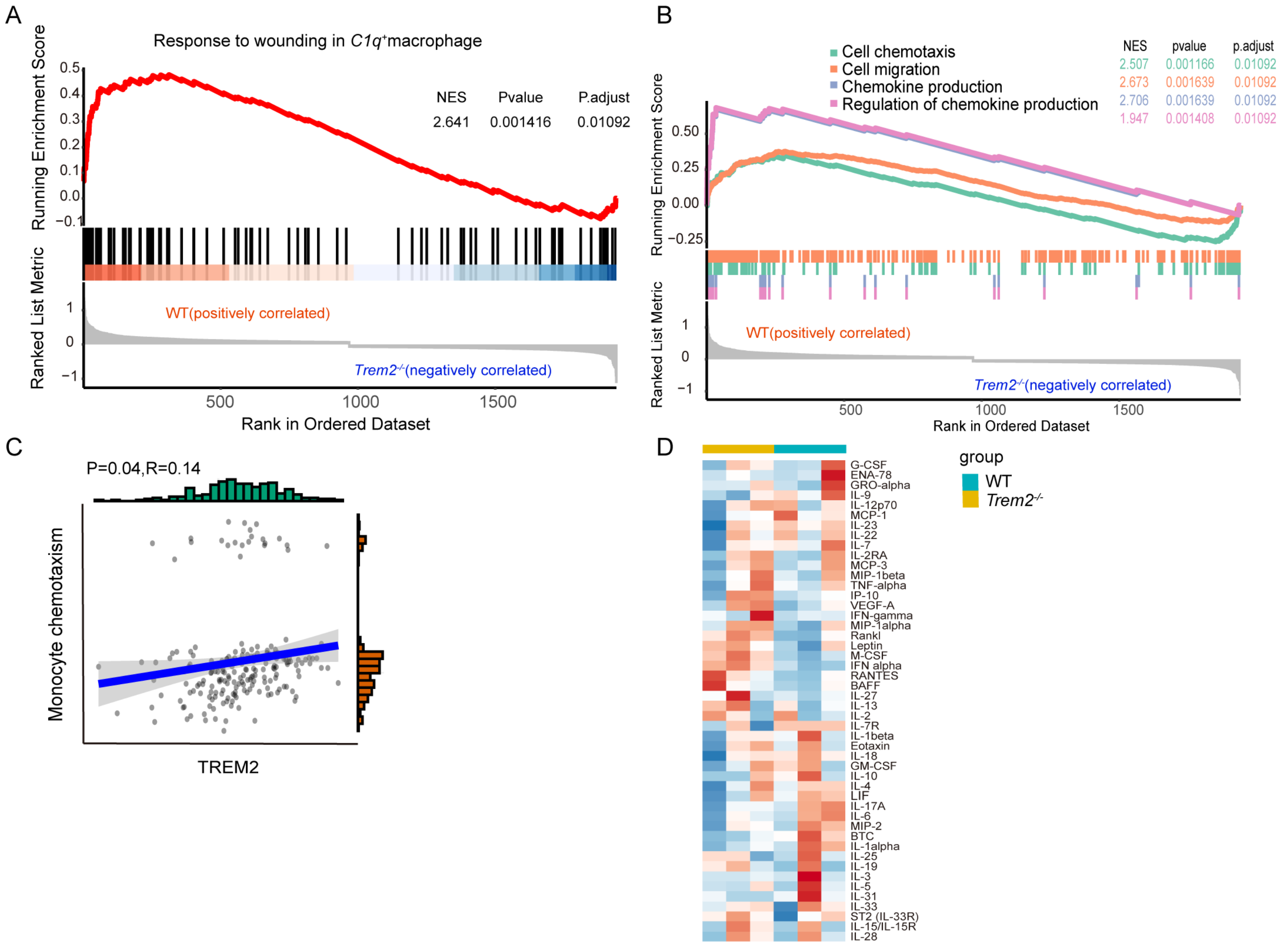
